# Localization microscopy at doubled precision with patterned illumination

**DOI:** 10.1101/554337

**Authors:** Jelmer Cnossen, Taylor Hinsdale, Rasmus Ø Thorsen, Florian Schueder, Ralf Jungmann, Carlas S. Smith, Bernd Rieger, Sjoerd Stallinga

## Abstract

MINFLUX offers a breakthrough in single molecule localization precision, but suffers from a tiny field-of-view and a lack of practical parallelism. Here, we combine centroid estimation and illumination pattern induced photon count variations in a conventional widefield imaging setup to extract position information over a typical micron sized field-of-view. We show a near twofold improvement in precision over standard localization with the same photon count on DNA-origami nano-structures.

## Main text

Single-molecule localization microscopy^1,2,3^ circumvents the diffraction limit using centroid estimation of sparsely activated, stochastically switching, single-molecule fluorescence images. Improvement over state-of-the-art image resolutions of around 20 nm towards values below 5 nm is desired for truly imaging at the molecular scale. This needs improvements in labelling strategy to reduce linker sizes^4,5,6,7^ and methods to overcome low labelling density such as data fusion^8^, but also a step in localization precision. Efforts so far have targeted an increase in the number of detected photons *N* by chemical engineering of brighter fluorophores^9^, or by avoiding photo-bleaching via e.g. cryogenic techniques^10,11,12^. These improvements scale the localization precision with 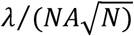, with *λ* the fluorescence emission wavelength, and *NA* the microscope objective numerical aperture^13^.

Recently, a new concept called MINFLUX was proposed^14^, in which a doughnut illumination spot is shifted over an area of size *L*~50 nm, and the position of a single molecule in the scan range is determined by triangulation based on the detected photon count for the different doughnut positions. The localization precision of this procedure scales as 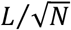, which is advantageous compared to 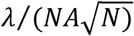, as the scan range *L* can in principle be chosen arbitrarily small. Drawbacks of MINFLUX are the limited field-of-view (FOV), and the low throughput, as the molecules are imaged one molecule at a time in the tiny Region Of Interest (ROI) of size *L*. Balzarotti et al. suggested the use of sinusoidal illumination patterns^14^, similar to those used in Structured Illumination Microscopy (SIM)^15^, and used earlier for single molecule tracking^16^. The extension of the triangulation procedure to spatially extended illumination patterns, however, remains a challenge.

Here, we propose to extract the molecule’s position in a combined estimation from *both* the relative position with respect to the shifting sinusoidal illumination pattern during all camera frames within the molecule’s on-event *and* from the estimated centers of the detected spots on the camera. This solves the challenge of photon count based localization with spatially extended illumination patterns. Our method, that we call SIMFLUX, overcomes the limited FOV and throughput of MINFLUX, and is compatible with standard widefield imaging on a camera. SIMFLUX is realized by a novel optical architecture for fast millisecond time scale switching of orthogonally oriented sinusoidal illumination patterns, and by a novel data processing strategy for spatiotemporal localization in relation to the shifting illumination patterns.

Figure 1a shows our optical architecture. A fast operable Pockels-cell switches between the two arms of a polarizing beam splitter in which piezo mounted gratings are placed that deliver the diffraction orders for interference based sinusoidal illumination patterns along two orthogonal directions (see Methods for details). This enables cycling through 6 patterns (2 orientations, 3 phase steps) on the millisecond time scale with sufficient power throughput. Only two orientations are needed, because this suffices for a Fisher-matrix that gives rise to an isotropic region of confidence for localization in the xy-plane (see Supplementary Note). This differs from SIM, where three or five orientations are needed for a near isotropic filling up of the support in image Fourier space^15^.

**Figure 1.**
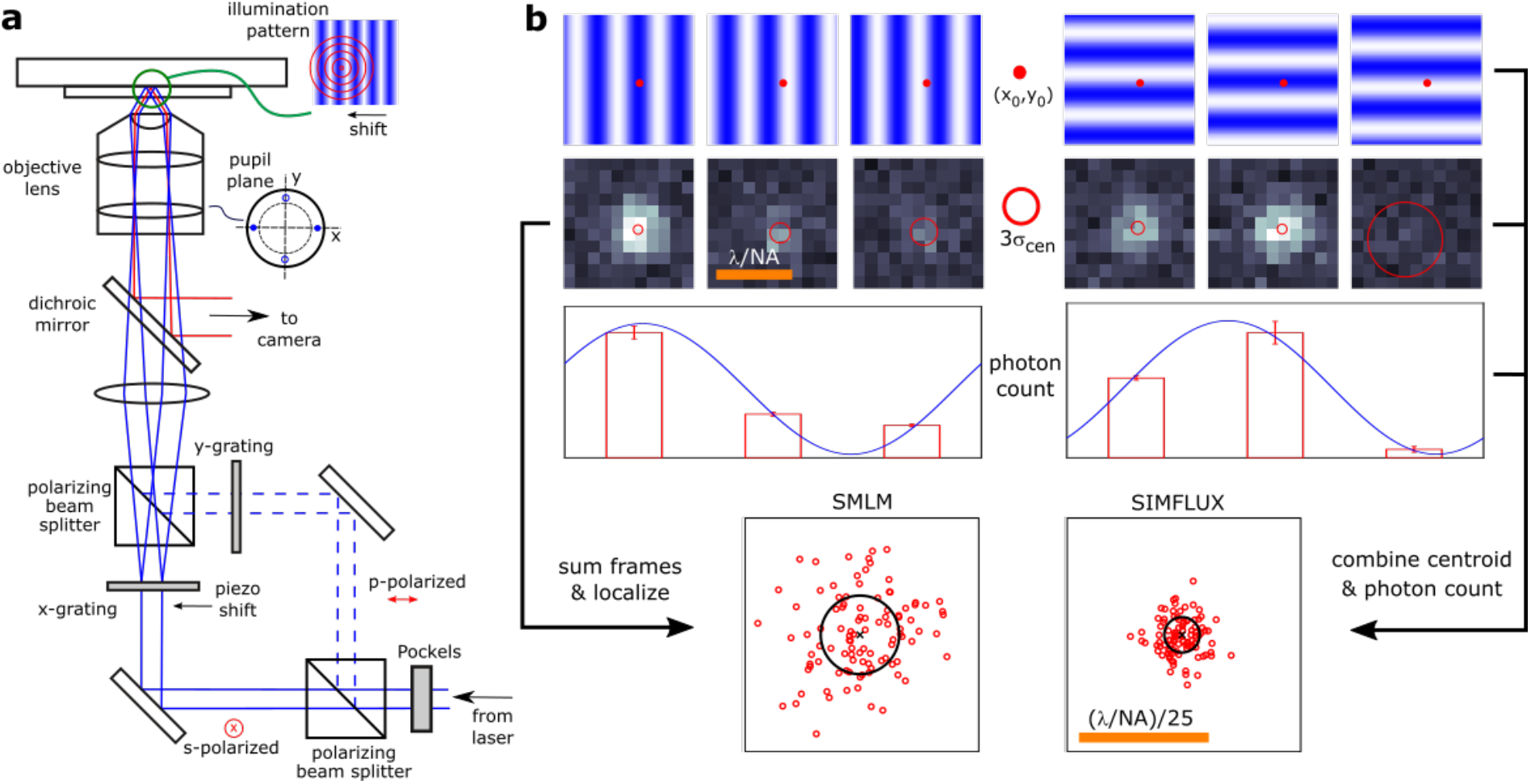
Principle of SIMFLUX. (**a**) A sinusoidal illumination pattern is created in a Total Internal Reflection (TIRF)-SIM setup by two counter propagating evanescent waves. Fast switching between two orthogonal line patterns is achieved by placing two piezo mounted gratings in the two arms of a polarizing beam splitter, selecting the operational arm by a polarization switching Pockels cell. (**b**) A total of 6 images are recorded with 3 shifted patterns per orthogonal orientation of the line pattern. Combining the centroid estimates of the 6 frames with the photon count in relation to the pattern shift improves the localization precision with a factor of around two compared to the standard centroid estimate on the sum of the 6 frames.

The processing pipeline (see Methods for details) requires the detection of single molecule emission events in space as well as in time, in combination with a retrieval of the illumination pattern parameters (pitch, orientation, modulation depth, and three phases per orientation, and relative intensity of the two beam splitter arms). First, the entire set of acquired images is processed using a standard SMLM pipeline for selecting ROIs per frame and for an initial localization fitting. This is done on the moving sum of 6 frames in order to enhance SNR for robust initial on-event detection. Next, the photon count is estimated for all individual frames within the 6 frame blocks, and blocks where the molecule is not in the on-state in all 6 frames are filtered out by a photon count threshold on the first and last of the 6 frames. Then, the pitch and orientations of the patterns are estimated using Fourier domain peak finding^17^ on the localization reconstruction. The pattern phases are subsequently retrieved by fitting the sinusoidal illumination pattern to the estimated single-frame photon counts. Next, the ROIs in the frames belonging to a molecular on-event are fitted with a Maximum Likelihood Estimation (MLE) routine, taking into account the centroid positions in each frame *and* the fluorescence signal strengths in relation to the shifting illumination pattern. The difference in the average position of these SIMFLUX localizations and the corresponding SMLM localizations is indicative for an error in the estimation of the pattern pitch and orientations, and can therefore be used to adjust the estimates. After updating them, a next round of pattern phase estimation and SIMFLUX fitting can start. This iterative procedure converges in 3-4 rounds.

The Cramér-Rao Lower Bound (CRLB) for the localization precision (see Supplementary Note) is given by:

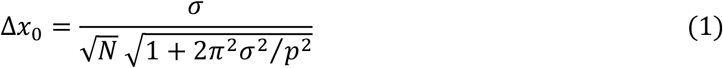

with *σ* ≈ *λ*/4*NA* the width of the Point Spread Function (PSF), and *N* the total number of collected photons during the on-event of the molecule. The smallest pitch of the standing wave illumination pattern is *p* ≈ *λ*/2*NA*, implying that the improvement factor over the SMLM precision 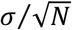 can reach values up to around 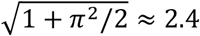. An imperfect modulation depth *m* (between 0.90 and 0.95) indicates a lower improvement factor of close to 2 (see Supplementary Note). Simulations with Gaussian and vector PSFs show that our method achieves the CRLB for a wide range of realistic photon counts and background photon levels (Supplementary Figs. 1 and 2). It appears further that background has the same relative impact as in conventional SMLM, implying that SIMFLUX can be used under the same experimental conditions as conventional SMLM^13^ (Supplementary Fig. 3). Simulations further show that to reach a twofold improvement in localization precision the modulation must be at least 0.9, and must be known with a precision of around 0.04, for the pattern phases a precision of ~2 deg is required (Supplementary Figs. 4 and 5). We meet these conditions in our experiments. Supplementary Fig. 6 shows that there are small variations in localization precision depending on the position of the molecule with respect to the minima of the illumination patterns, leading to improvement factors compared to conventional SMLM that range between 1.6 and 2.3, with an average of 2.1 (for *p/σ* ≈ 2 and *m* ≈ 0.95). These variations are reduced by increasing the number of phase steps (Supplementary Fig. 7 and Supplementary Note).

We have tested our method on DNA-origami nano-structures imaged with DNA-PAINT^18^ (see Methods). Figure 2a shows the SIMFLUX reconstruction over the full 26×26 μm FOV of nano-rulers with binding site spacing of 80 nm, Fig. 2b-d show 5 selected SIMFLUX nanoruler images across the FOV, with improved precision compared to the SMLM images. The latter uses the fits from the sum of 6 frames used for SIMFLUX, which effectively provides a spatially uniform illumination single-molecule image. The projections of the localizations in Figs. 2d,e on the *x*-axis provides localization histograms (Figs. 2f,g), indicating an improvement in localization precision with a factor of around 2. The localization precision, measured from the accumulated data of 432 segmented binding sites across the whole FOV, improves from 16.8 nm to 9.4 nm (Figs. 2h-j), an improvement factor of 1.8, and is comparable to FRC-resolution values^19^ 15.5 nm and 8.4 nm (Supplementary Fig. 9), an improvement factor of 1.9. The achieved precision does not match the CRLB (Supplementary Fig. 10), which we attribute to a residual drift of around 4 nm. This level of residual drift after correction is reasonable in view of the sparsity of the sample. Drift may also be the root cause for the washing out of the dependence of the precision on global phase, anticipated by theory, and for a SIMFLUX precision improvement factor that is somewhat less than the theoretical value 2.1 (for *σ* = 119 nm, *p* = 220 nm, *m* = 0.92). A similar precision improvement of 2.0 can be achieved for the case of 4 phase steps (Supplementary Figure 11), which can provide more robustness against errors in detecting the on-off transitions. Figs. 2k-o and Supplementary Fig. 12 show further results on DNA-origami grids with binding site spacing of 40 nm and 20 nm, showing similar resolution improvements.

**Figure 2.**
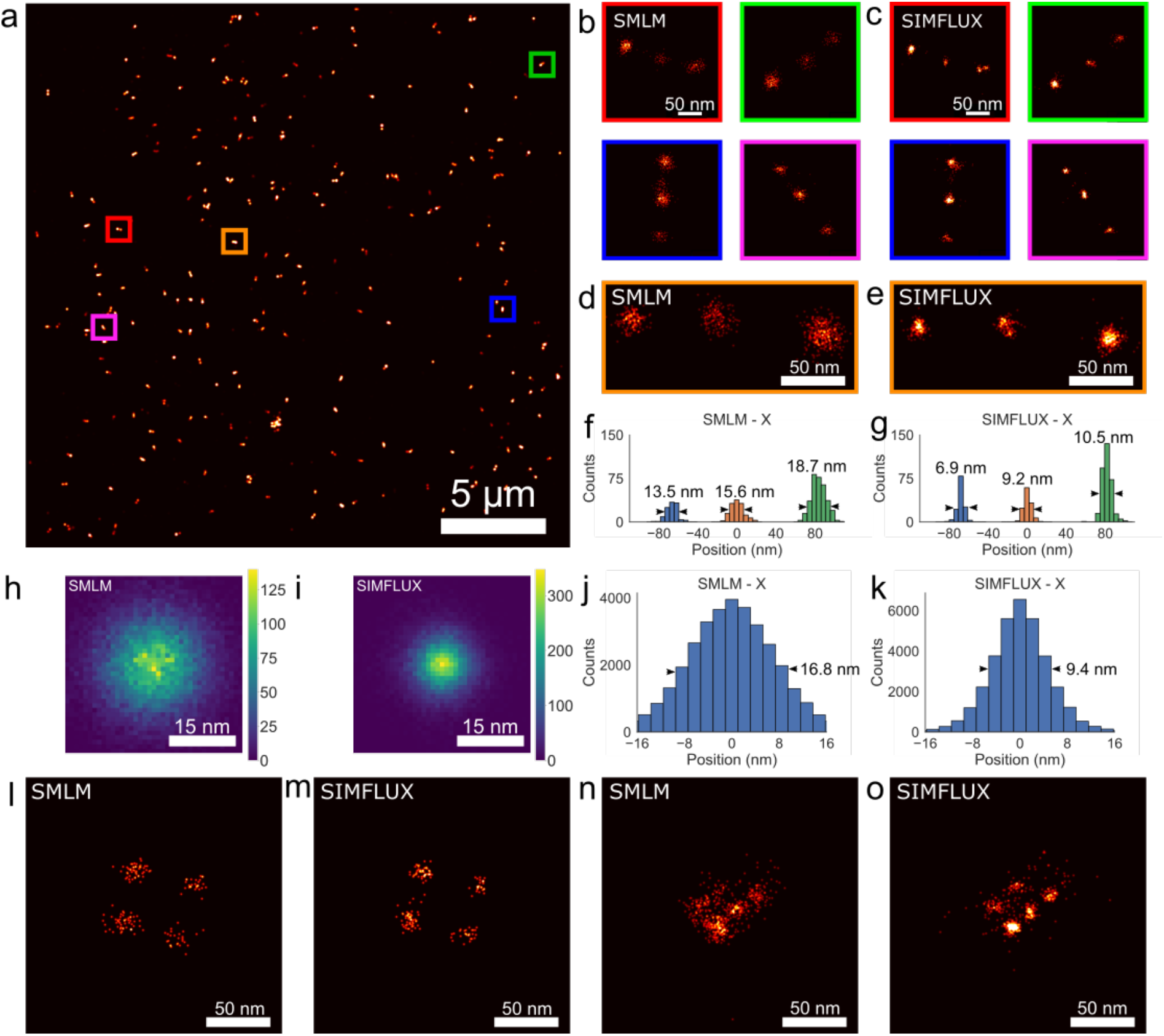
Demonstration of SIMFLUX on DNA-origami nano-stuctures. **(a)** Full 26 μm wide FOV SIMFLUX image of sparsely distributed nanorulers with 80 nm spacing. **(b,c)** Zoom-in on 4 conventional SMLM and SIMFLUX nano-ruler instances color indicated as boxes in (a). **(d,e)** SMLM and SIMFLUX image of nanoruler instance of box in (a). **(f,g)** Histograms of localizations in (c,d) projected on the *x*-axis, showing a near twofold improvement in precision. **(h,i)** 2D histograms of SMLM and SIMFLUX localizations in the image plane, assembled from 432 segmented binding sites, and **(j,k)** histograms of localizations projected onto the *x*-direction, showing an isotropic improvement in measured FWHM of 1.8. **(l-m)** SMLM and SIMFLUX images of DNA-origami grids with 40 nm spacing between binding sites, showing a clear gain in precision of SIMFLUX. **(n,o)** SMLM and SIMFLUX images of DNA-origami grids with 20 nm spacing, hardly resolved in SMLM, clearly in SIMFLUX.

In conclusion, we have demonstrated a practical way to extend the MINFLUX concept to sinusoidal illumination patterns, improving field-of-view and throughput to standard SMLM experimental settings. We envision that our technique can also be used to achieve the same precision as SMLM but with fourfold less light, enabling either faster imaging or imaging with dimmer fluorophores. Our optical setup can in principle achieve the same resolution gain as MINFLUX over a small FOV of size *L*, much smaller than the pattern pitch *p*, if we shift dark fringes of the pattern over a total translation range *L* instead of *p* (see Supplementary Note, and Supplementary Fig. 13). We also envision advances in data processing by application of e.g. hidden Markov-models to precisely model the on-off transitions. Another next step for SIMFLUX would be the extension to 3D interference patterns for an improvement in both lateral and axial localization precision.

## Author contributions

Imaging experiments were done by T.H. and F.S., data analyses were done by J.C. and T.H., simulations were done by R.Ø.T. and J.C., F.S. and R.J. provided samples. C.S.S., B.R., and S.S. devised key concepts, supervised the study and wrote the paper. All authors read and approved the manuscript.

## Acknowledgements

J.C. and C.S.S. were supported by The Netherlands Organisation for Scientific Research (NWO), under NWO START-UP project 740.018.015 and NWO Veni project 16761, T.H., R.Ø.T. and B.R. acknowledge National Institutes of Health (Grant No. U01EB021238), F.S. and R.J. acknowledge support by an ERC Starting Grant (MolMap, grant agreement no. 680241).

## Data and code availability

Data and code will be made available upon publication with a doi-identifier (data) and at github (code).

## Methods

### Experimental setup

A custom total internal reflection (TIRF) structured illumination microscopy (SIM) microscope was built to implement the SIMFLUX method (Supplementary Figure 14). The setup uses a 200 mW, 640 nm, diode laser (Toptica, CLUP-640) that is spectrally filtered with a 640/20 nm (Chroma, ET640/20m) bandpass filter and spatially filtered by coupling into a polarization maintaining single mode fiber (ThorLabs, PM630-HP) via an NA matched aspheric lens, L1 (f = 3.3 mm, ThorLabs, C340TMD-A). The output of the fiber is collimated by an objective, L2 (0.45/20X A-PLAN, Zeiss). SIMFLUX utilizes two orthogonal sinusoidal modulation patterns in the focal plane of the objective lens. The optical architecture overcomes drawbacks of typical SIM architectures. Rotating gratings are too slow to generate multiple illumination patterns for a typical molecular on-event (~10s of ms), Spatial Light Modulators (SLMs) are sufficiently fast, but too power-inefficient to generate a sufficiently high illumination intensity (~kW/cm^2^) over an extended field of view (~10s of μm). A simple way to generate these is to build an interferometer and self-interfere a laser at the sample plane. To do this, custom etched binary phase gratings (HOLOOR, DS-281-1-Y-A) with pitches of 8.496 μm are used to generate ±1st diffraction orders with near theoretical diffraction efficiency limits of around 79%. Distinct and orthogonal interference patterns at the focal plane with controllable phase are generated using a fluid filled KD*P Pockels cell (Leysop, EM508-2T-F-AR640) to alternate the laser between two beam paths and piezoelectric stages (PI, P-753.1CD) to phase shift the binary phase gratings. Before being sent through the Pockels cell and diffraction gratings, the laser intensity is controlled via a half wave plate (ThorLabs, DS-281-1-Y-A) and a Glan-Taylor polarizer (GL10-A) to attenuate when needed while maintaining at least a 1000:1 intensity extinction ratio between each path. The beam then passes through the Pockels cell that is aligned such that applying a half wave voltage switches the beam between s and p polarizations. Two mirrors (ThorLabs, PF10-03-G01) then align the laser to the main optical axis of the system. A quarter wave plate and half wave plate (ThorLabs, WPQ05M-633 & WPH05M-633) are placed after the second mirror to reduce any elliptical polarization induced by reflection. A cube polarizing beam splitter (ThorLabs, CCM1-PBS252/M) selects the beam path based on s or p polarization entry. A high extinction ratio Glan-Taylor polarizer (ThorLabs, GL10-A) is then placed in each beam path after the polarizing beam splitter to ensure at least 10^4^ polarization purity in each beam path. A binary phase grating is then placed in both beam paths. Each grating is mounted on a nanometer resolution piezoelectric translation stage to induce phase shifting. The gratings are aligned on the piezoelectric stages so that their main diffraction axes are orthogonal to one another. The azimuthal alignment of the gratings is chosen such that the polarization of the interfering diffraction orders is parallel in the objective focal plane for each beam path. After light is diffracted from each binary phase grating, a second polarizing beam splitter recombines the two paths into the main system illumination path. Two more beam steering mirrors are needed to recombine the beam path that is reflected off the first beam splitter. After recombining into a single optical axis, the diffracted orders are collimated by L3 (ThorLabs, ACA254-075-A) and sent through a spatial filter mask to filter all but the ±1st diffraction orders. From there a 4f system L4,5 (Edmund Optics/ThorLabs, 49-395-INK/AC508-180-A-ML) relays the spatial filter to the rear focal plane of the objective (Nikon, CFI Apo 1.49 TIRF 100XC Oil) after reflecting off a long pass dichroic mirror (Semrock, Di03-R660-t1-25.2×35.6). If the light from the ±1st orders is well focused in the rear focal plane, collimated light will emerge from the objective and be incident on the sample plane. The ±1st orders enter at opposite edges of the back focal plane at a radius ±2.91 mm from the optical axis, corresponding to a Numerical Aperture *NA_i_* = 2.91/2.0 = 1.455 (the focal length of the Nikon 100x objective lens is 2.0 mm). The illumination *NA_i_* exceeds the sample refractive index of 1.33 and therefore provides TIRF illumination. A TIRF illumination system is chosen in order to reduce background fluorescence by providing an interface bound optical sectioning of 100-200 nm, and to be compatible with DNA-PAINT based localization. Control of the sample plane and system focus is achieved with a XYZ 100×100×100 μm travel range piezoelectric slide stage (Mad City Labs, 1D100). Emitted fluorescence is collected by the same Nikon objective in an epi-illumination configuration and passes through the long pass dichroic mirror and a bandpass 690/50 nm emission filter (Chroma, ET690/50m) before being imaged by an infinity corrected tube lens (ThorLabs, TTL200-A) onto an sCMOS camera (Hamamatsu, ORCA Flash 4.0 V2). The pixel size of our camera in the sensor plane is 6.5 μm giving a back-projected pixel size in the sample plane equal to 65 nm. Image acquisition was controlled using a standard desktop workstation equipped with a camera link frame grabber (Hamamatsu, AS-FBD-1XCLD-2PE8). Micro-manager serves as the main image acquisition software, but it is integrated with a custom Python script to control an Arduino which triggers the PI piezoelectric stage controllers and the Pockels cell to iterate through imaging states. Micromanager also controls the piezoelectric sample stage from Mad City Labs. The PI piezo electric stages were initialized to receive triggers from the Arduino via the program MikroMove.

### Calibration of modulation contrast

The modulation contrast of the system was characterized by directly probing the sinusoidal pattern. This was done by imaging a sparse single molecule sample and finely phase shifting the illumination pattern over the sample by supplying the piezos with incremental voltage steps. By imaging after each phase shift, a direct measurement of the sinusoidal wave can be traced over the image series duration. Our samples were pre-prepared slides of 20 nm GATTA-beads (GattaQuant, Bead R). Regions of interest that contain only a single bead, of size 11×11 pixels, can be extracted from the image series and the ADU, or total photons if converted, can be calculated in each frame. A sinusoidal curve *y* = 1 + *m* sin(2*πx*/*p* + *ψ*) can be fit to this data with an *R*^2^ > 0.98 and the modulation contrast *m* can be extracted. Our images have a small amount of scattered background fluorescent signal that rises above the camera offset of 100 ADU. The typical fluorescent background of a sparsely populated sample in our TIRF illumination microscope is 105 ADU on average. Photo bleaching of the fluorescent beads not only changes the spot intensity, it also makes the scattered fluorescence caused by non-TIRF illumination decrease. Consequently, the background changes slightly over the duration of the image series. This changing background fluorescence was estimated by fitting a linear decay function. The line was then used to estimate the time dependent background to be subtracted for each given frame.

Supplementary Figure 15 shows an example of a segmented 11×11 pixel ROI at three different points along the modulation pattern. The three ROI images were taken from a larger 200 image series where more than a full period of the modulation pattern was swept across the bead. To quantify the modulation contrast for the full field of view, many beads (198 in total) were localized, their ROIs extracted, and their summed ADUs fitted with a sinusoid as seen in Supplementary Figure 15. The distribution of fitted modulation contrast values in the two pattern directions shown in Supplementary Figure 15 have median values of 0.91 and 0.92, respectively. However, there are outliers that tail off towards low contrast values. Some of these spots may not be effective point sources, e.g. beads that have other particles in close proximity, false segmentations where a region of interest is largely noise, or regions of interest with more than one bead. This would adversely affect the measurement of the modulation contrast. These could be omitted as outliers for this calibration, but it is not necessary. For example, excluding points with *m* < 0.75, has no effect on the median *k_y_* modulation contrast and only raises *k_x_* modulation contrast by 0.01. Approximately 80-90% of the segmented calibration beads fall into the main body of the histogram (*m* > 0.75) where the total amount of analysed ROIs in the full histogram amounts to 184.

### Initial estimate of pitch and phase to piezo voltage calibration

The pitch of the interference pattern in the sample plane can be calibrated by using high density, blinking, fluorophores that are evenly distributed in the sample plane. The interference pattern is realized by localizing the fluorophores under the standing wave illumination. Normally, the pattern is diffraction limited and not visible directly, however, super-resolved localization images show it quite clearly. Taking the Fourier transform of the image allows for direct measurement of the pattern spatial frequencies, equal to 219.94 nm, as seen in Supplementary Figure 16. The initial estimate of the pitch agrees well with the expected value *λ_ex_*/(2*NA_i_*) = 640/(2 ∗ 1.455) = 219.9 nm. This value is used as the initial estimate for the spatial frequency vector estimate in the data analysis.

The direct calibration of stage translation to phase for the piezo mounted diffraction gratings, both relative and absolute, can be calculated from this data as well. Each frame from the sinusoidal fit data in Supplementary Figure 15 is separated by steps of 0.04 μm at the piezoelectric plane. The median fit period is found to be 106.14 frames. This gives 1.42 μm per third of a period. This can then be conveyed in the sample plane as 51.6 nm/μm, where the μm refer to the translation distance in the grating plane.

### Samples

Gattaquant nanorulers based on DNA-PAINT, GATTA-PAINT (PAINT 80R ATTO 655), were used as the main samples for our imaging experiments. They consist of three equally-spaced binding sites separated by 80 nm between each with an approximate surface density of 1/μm^2^. Other DNA-PAINT based nanostructures were imaged with uniformly decreasing structure sizes: 2×2 grids with 40 nm binding site distance and 4×3 grids with 20 nm binding site distance (see Supplementary Fig. 16 for designs) were synthesized and prepared according to the protocols provided by Schnitzbauer et al.^18^ employing 5’-TTATACATCTA-3’ as DNA-PAINT docking strand (positions marked in red in Supplementary Fig. 17) and 5’-CTAGATGTAT-3’-Cy3B as DNA-PAINT imager sequence. Both nanostructures were imaged using 5 nM imager strand concentration.

### Data acquisition

A simple data acquisition sequence was defined to acquire six (or any other arbitrary number) phase shifted images during the on-time of a single blinking event of a fluorophore (Supplementary Figure 18). Here, the digital trigger for the acquisition is a digital TTL generator that triggers a frame capture event on the sCMOS camera. The camera outputs a high TTL when all pixels experience a global exposure, e.g. all pixels will integrate photons for the same amount of time. The falling edge of the global exposure sequence triggers an Arduino to cycle through a three bit digital output that controls the two translation piezos and the Pockels cell. The Pockels cell alternates the laser beam between s and p polarization states to determine which diffraction grating is illuminated. This event is triggered by the Arduino such that the polarization switching occurs between periods of global pixel exposure. Likewise, the two piezos which hold the diffraction gratings are triggered to move by the Arduino. Each piezo mounted grating moves while the opposite grating is illuminated and imaged, thus maximizing imaging speed. To ensure that illumination from the s polarized arm does not appear in images from the p polarized arm, and vice versa, the laser was pulsed to only illuminate during the camera global exposure. For the samples in this experiment, the average on-time of blinking events is ~200-300 ms for the GattaQuant nano-rulers, ~1 s for the 40 nm 2×2 grids, and ~100 ms for the 20 nm 4×3 grids. All samples are imaged at 70 frames per second, ensuring that a full phase cycle of each pattern orientation is captured. Future experiments could be imaged at much higher frame rates. Typical fluorophore on times for STORM and PALM can range from 10-60 ms. At 70 frames per second, a fluorophore with an average on time of 10-60 ms would not be present in all 6 frames, three for each pattern orientation, required for SIMFLUX. Using our current sCMOS, full sensor readout takes approximately 10 ms, limiting the full field of view frame rate to 100 frames per second. Vertical cropping linearly reduces the amount of time required to read out the frame, increasing the potential frame rate. Our raw image region of interest is 400×400 pixels in the center of the sCMOS field of view and has a theoretical frame rate of 512 frames per second, or 1.9 ms per frame. Practical limitations stopped us from imaging at this high of a frame rate. The piezoelectric stages that the gratings are mounted on have a step and settle time of 3-4 ms. Using an exposure time of 3-4 ms, even with our alternating data acquisition scheme, would likely result in the piezoelectric stages moving or settling to their final positions during an exposure. Still, a conservative 200-300 frames per second, around 20-30 ms per six images, would be enough for STORM and PALM type imaging and plausible in future implementations. Laser power density at the sample is the second consideration for fast imaging. Our current system illuminates the sample plane with a power density of ~600 W/cm^2^ over an 80 μm illumination diameter. However, only 26 μms in the center of the illumination are used to maintain a relatively uniform excitation intensity profile. This is generally too low for STORM, requiring higher power densities on the order of 2-4 kW/cm^2^. However, DNA-PAINT does not require such high power densities and 600 W/cm^2^ was sufficient for imaging at 70 frames per second. Future iterations can incorporate a high power laser, increasing our initial laser power from 200 mW to 1 W and the power density at the sample from 600 W/cm^2^ to 3 kW/cm^2^.

### Simulation setup

Simulated point spread functions (PSFs) are generated according to a vectorial PSF model^20^. The NA is taken to be 1.49, the wavelength 680 nm, the refractive index 1.515 (medium, cover slip and immersion fluid assumed to be matched), with a pixel size of 65 nm in object space, and the region of interest (ROI) is 11×11 pixels large. The PSF coordinates within the ROI are drawn from a uniform distribution with a width of half the illumination pattern pitch. Unless stated otherwise, we take 6000 detected signal photons and 30 background photons per pixel, and we add noise according to Poisson statistics. The simulation is run for 5000 randomized instances. The excitation pattern is taken to be sinusoidal with three patterns along the *x* and y-axis. The pitch is taken to be 243.75 nm (3.8 pixels), which is set equal to about 2× the spot width for the sake of simplicity. The number of signal photons reported corresponds to the number of photons captured over the entire FOV, i.e. taking into account the spatially extended tail of the PSF that falls outside the ROI^21^.

We have also used simulations of blinking emitters over a full FOV. A series 12,000 images is simulated over a FOV of size 256×256 pixels^2^ with a pixel size of 65 nm. The imaged structure is a 24×24 grid of binding sites with a spacing of 650 nm. The actual emitter coordinates are drawn from a uniform distribution with a width of 110 nm around the binding sites, introducing randomness to the structure while keeping the inter-emitter distance sufficiently high for having well-separable spots in the simulated images. The illumination pattern is shifted in 3 steps over the pitch of 220 nm with a modulation depth of 0.95 in both the *x* and *y*-direction to match the expected experimental values. Blinking events are generated for each emitter using the transition rates k_on_ and k_off_ for dark to active, and active to dark transitions respectively. The rates were fixed at k_on_= 0.001 frame^−1^ and k_off_= 0.04 frame^−1^, giving a typical on-time of 25 frames and a typical off-times of 1000 frames. Random transitions between both states were simulated at a rate of 3× the frame-rate. The locations of the resulting set of emitters that are are in the on-state in a frame are blurred with the vectorial PSFs as described above. Shot noise is subsequently added, using, unless stated otherwise, 2000 detected signal photons per spot and 10 background photons per pixel.

### Processing pipeline

Supplementary Figure 20 gives a schematic overview of the entire processing pipeline. First, the set of acquired images where first offset and gain corrected to convert ADUs into photons^22,23^. The total set of acquired images is 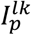 with *l* = 1,…, *L* the pattern orientations, *k* = 1,…, *K* the pattern phases, and *p* = 1,…, *P* the label for the groups of *L* × *K* frames, giving a total of *L* × *K* × *P* acquired frames. The detection of isolated emitting molecules is aided by first applying a sum over the *L* × *K* blocks of frames, i.e. the set of 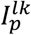 is summed to 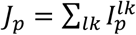. This averages out the effect of the shifting and rotating illumination pattern, and increases the Signal-to-Noise Ratio (SNR) for spot detection. ROIs of size 9×9 pixels are identified by a two-stage filtering process to reduce photon noise and local background followed by an intensity threshold ^24,25^. In short, we apply uniform filters to the raw images with filter size 4 and 8 pixels and take the difference. We then computed the local maximum in a 5×5 pixels neighbourhood for all pixels and accept the central pixel as candidate for a single-molecule spot if its value is the local maximum and is higher than a threshold of 10 (for the nanoruler dataset of Fig. 2 and Supplementary Fig. 11) or 20 (for the grid DNA-origami datasets of Fig. 2 and Supplementary Fig. 12). Now a 9×9 pixel ROI is segmented out for all candidates and each ROI, labelled with index *s*, is extracted and fitted for emitter position 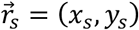, signal photon count *N_s_* and background *b_s_* using established Maximum Likelihood Estimation (MLE) fitting^26^ using a Gaussian PSF model. The fits are done with a fixed Gaussian spot width of 119 nm, determined from a separate fit on the first few frames of the entire dataset.

In a next step the signal photon count and background in the ROI with label *s* in the *L* × *K* original individual frames 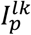 are analysed for estimating the signal photon count 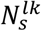 and background 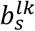 given the estimate of the emitter position (*x_s_, y_s_*) obtained from fitting the moving sum images *J_p_* (see Supplementary Note for details). By keeping the position fixed a more robust estimate of photon count can be made in frames with low photon count, intermittent in a series of frames where the molecule is brighter. This can occur when a dark fringe of the illumination pattern passes by at the molecule’s position. It is mentioned that using a Gaussian PSF model gives rise to an underestimation of the signal photon count^21^ by ~30%. In the end we only use the information from the relative signal photon count for different phases and orientations of the illumination pattern for fitting the position of the molecule, implying that the same underestimation for all photon count estimates has a limited impact on the final localization precision. Simulations confirm that Gaussian PSF fitting works reliably for realistic signal photon counts (up to about 10^4^), with a need for more sophisticated PSF models only arising for extremely bright emitters (see Supplementary Figs. 1 and 2).

Sequences of *L* × *K* frames where the molecule is in the off-state in the first few or last few frames are rejected by application of a minimum filter. Molecular on-events are selected where the first and last frames are above a user set minimum number of photons, i.e. 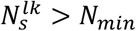 is required for the *l* and *k* corresponding to the first or last of the block of *L* × *K* frames. The choice for the threshold *N_min_* is based on a simulation study (see Supplementary Fig. 19 for the key results). It appears that for our experimental parameters (around 1500 signal photons and 10 background photons/pixel) the setting *N_min_* = 30 results in a near-zero false positive rate and a false negative rate around 20%.

After application of the minimum filter we proceed with estimating the illumination pattern parameters from the data. First, we make an initial estimate of the spatial frequency vectors 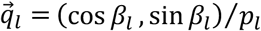 (with pitch *p_l_* and orientation *β_l_*) of the patterns. The set of molecular on-events with label *s* contains *L* × *K* single-frame localizations with estimated coordinates (*x_s_, y_s_*), signal count 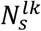 and background 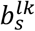. The entire collection of these single-frame localizations is split into subsets corresponding to the *l* = 1,.., *L* orientations and *k* = 1,…, *K* phases of the illumination patterns. The *L* × *K* subsets of single-frame localizations are used to generate super-resolution reconstructions 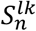 defined on a grid of super-resolution pixels 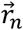, with *n* the index of the super-resolution pixels. We have used Gaussian blob rendering with a width equal to the average localization precision from the single-frame localizations, and a zoom factor of 6 compared to the detector pixel grid to make the superresolution pixel size comparable to the single-frame localization precision^19^. For the data of Fig. 2 we have used a super-resolution pixel size equal to 10.8 nm, comparable to the CRLB in the single-frame localizations of around 12 nm. Each Gaussian blob is multiplied with a weight factor equal to the estimated signal photon count 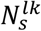. The spatial frequencies 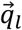 are then detected by finding the peak in the Fourier domain of the reconstructions 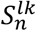 (see also Supplementary Fig. 20), improving over the initial estimate of the spatial frequencies 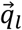, provided by the Fourier domain peak locations of the experimental calibration shown in Supplementary Fig. 10.

It appears that this first estimate of pitch and orientation of the patterns is not precise enough to obtain SIMFLUX localizations that have no position biases. This is solved by an iterative refinement procedure. The first step here is to estimate the illumination pattern phases *ψ_lk_*, as well as the modulation depths *m_l_*, and relative intensity *η_l_* for illumination patterns with orientation *l* (normalized as 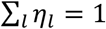, nominally *η_l_* = 1/*L*). These estimates are obtained by a least squares fit of the illumination pattern to the detected photon counts 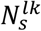 with error metric:

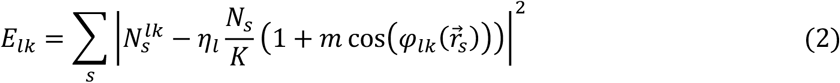

with 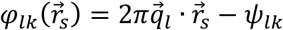 the phase at localization position 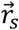. Illumination pattern phase estimation biases originating from the structure of the underlying fluorescently labelled structure are mitigated by taking into account the sum of all detected photon counts 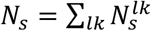 as weight factor for the illumination pattern in the error metric. The minimization of Equation 2 with respect to the 0th and 1st order Fourier coefficients (*η_l_, η_l_m_l_* cos *ψ_lk_, η_l_m_l_* sin *ψ_lk_*) of the sinusoidal illumination function results in the following set of linear equations:

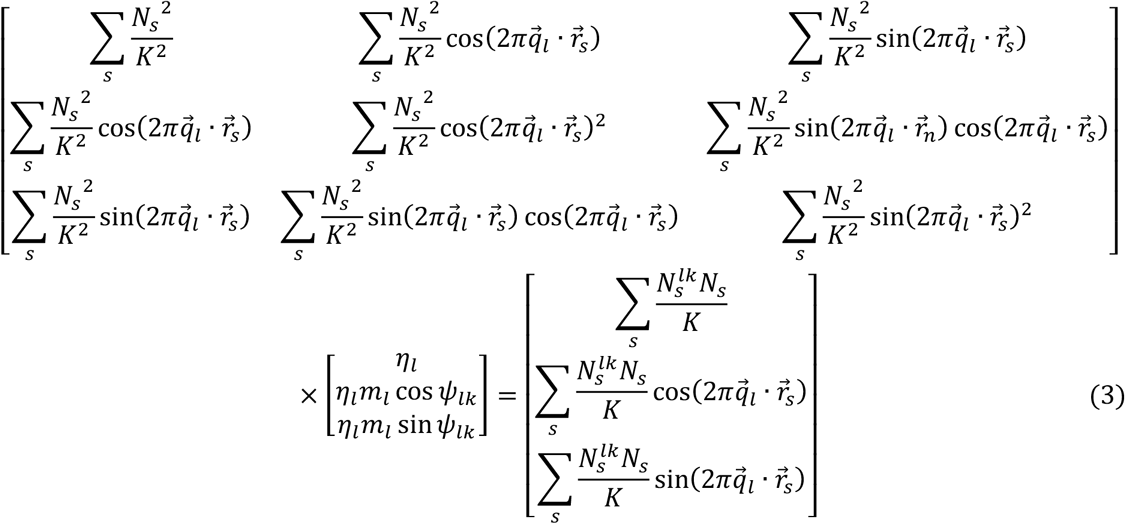

which can be solved in a straightforward way. In order to further enhance robustness of the fit an iterative procedure is applied in which the median of the quadratic error distribution over the localizations in Equation 2 is determined, and the localizations with error less than the median are kept for a second phase estimation. After this second phase estimation the median of the quadratic error of the original set of localizations is determined again, and the localizations with error less than the median are kept for a third phase estimation, etc. This procedure converges within 3 iterations. We apply this procedure on the set of localizations that is obtained before application of the minimum filter. In this way blocks of frames in which the molecule is partially in the on-state (say in the last 3 but not in the first 3 frames) aid in the fitting. The frames where the molecule is in the on-state provide genuine data points (the last 3 frames in the example), while the frames in which the molecule is in the off-state (the first 3 frames in the example) have no impact on the fit. The phase estimation finally has a standard error of the mean typically between 0.5 and 1.0 deg (see Supplementary Fig. 8), which we determine by splitting the entire dataset in 10 bins, repeating the phase estimation for the 10 bins, and computing the standard deviation over the estimated phases (normalized by 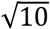). The modulation depths *m_l_* are typically estimated around 0.95, in agreement with the calibration measurements on beads. The modulation depth is typically underestimated for non-sparse datasets. In that case it is better kept fixed to 0.95, the typical value obtained for sparse datasets. The relative intensity *η*_1_ = 1 − *η*_2_ is found to be around 0.455 in our setup.

Next, an MLE based estimate is made of the molecule’s position, using both image centroid information and photon count information. The PSF model, log-likelihood, and relevant derivatives with respect to the fit parameters are defined in the Supplementary Note. Initial values for the parameter estimation are taken from the analyses on single-frame and moving sum frame data, the optimization uses the Levenberg-Marquardt algorithm. The previously estimated illumination pattern parameters are assumed to be constant throughout the experiment.

This SIMFLUX estimate differs 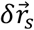 with the corresponding SMLM localization, where *s* labels the different localization events. An improved estimate of the spatial frequencies can now be made by minimizing the overall error in the illumination pattern phases 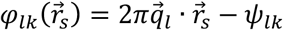. The average phase error per orientation is:

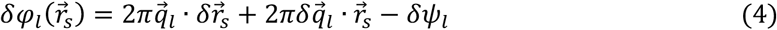

where 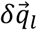 is the error in the spatial frequency vector, and where *δψ_l_* is the average error in the pattern phase. These errors can be estimated by linear regression, i.e. by minimizing:

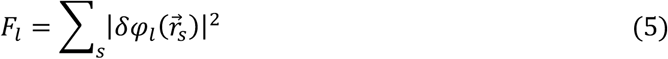

This results in a linear set of equations for 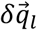 and 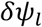:

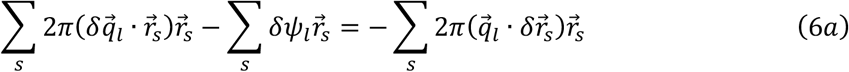

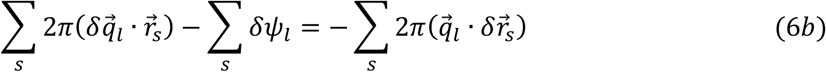

which can be solved in a straightforward way. After updating the spatial frequency vectors to 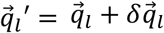 the estimation of the pattern phases *ψ_lk_* as explained above is repeated, as well as the SIMFLUX MLE fit. This procedure converges in 3 to 4 iterations.

The quality of convergence can be assessed by the rms value of the SMLM-SIMFLUX localization difference 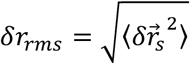. It appears that at convergence this rms value is about 13.0 nm for the nanoruler dataset of Fig. 2 (see Supplementary Fig. 20). This value is on the order of the localization uncertainty, which seems physically reasonable. It implies an error in the overall pattern phase of about 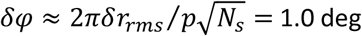 with *N_s_* = 431 the number of imaged binding sites used in the analysis and *p* = 220 nm the nominal pitch. This can be related to the final precision in the pitch estimation *δp*, which scales with the precision of the overall pattern phase estimation according to 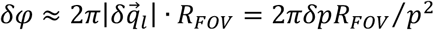, with *R_FOV_* = 13 μm the FOV size. Filling in the numbers then gives a precision in the pitch estimation of about *δp* ≈ 0.01 nm.

### Drift correction and data analysis

Sample drift is corrected on the localization data following the method of Schnitzbauer et al.^18^, appropriate for the sparse DNA-origami samples of our experiments. For this we have used the Picasso software tool, available at https://github.com/jungmannlab/picasso. In short, first a cross-correlation approach is used for a coarse drift correction. Then, multiple regions of interest are selected that contain only a single DNA-PAINT binding site. Typically, 20-30 binding sites are initially selected by hand with a diameter between 0.5 and 1.2 camera pixels. Each region of interest is selected such that there is both SMLM and SIMFLUX localizations present. The initial regions of interest are then used to automatically select similar binding sites based on the size of the region of interest and density of points within each one. This results in 400-700 binding sites when using the 80 nm nano-rulers from. After similar regions of interest are selected, a global drift correction is calculated and applied by minimizing the RMS deviation to the center of mass in each region of interest. We note that sample drift does not influence the pattern parameter estimation as the projected pattern is static under sample drift. Therefore, we do not need to re-estimate the pattern parameters after drift correction is applied to the localizations.

After drift correction, both the SMLM and SIMFLUX localizations in the selected regions of interest are exported and used to create the histograms for characterizing the spread of localizations. The localization point clouds are first mean shifted to zero and then added. A kernel density estimate of the histograms is used to measure the FWHM of the histograms. Further estimates of the spread of localizations that are computed are the Fourier Ring Correlation (FRC)^19^ of the entire super-resolution reconstructions (see Supplementary Fig. 9), and the standard deviation of the point clouds (see Supplementary Fig. 10).

All images are rendered by histogram binning on a grid with 0.52 nm (Fig. 2d,e,l,m,n,o and Supplementary Fig. 12) or 0.65 nm (Fig. 2b,c) super-resolution pixel size with additional Gaussian blurring with kernel size equal to 1 super-resolution pixel. The overview image Fig. 2a is rendered with a super-resolution pixel size of 34 nm.

## List of Supplementary Figures

1. Impact of signal photon count on the localization precision of SIMFLUX in comparison to standard SMLM for different background levels

2. Impact of signal photon count on the photon count estimation of SIMFLUX in comparison to standard SMLM

3. Impact of background photon count on localization precision of SIMFLUX in comparison to standard SMLM

4. Assessment of illumination pattern modulation estimation on localization precision in SIMFLUX

5. Impact of illumination pattern phase errors on localization precision and bias of SIMFLUX

6. Impact of emitter position in the global phase pattern on localization precision in SIMFLUX

7. Impact of modulation depth and global phase on zero-background CRLB of the localization precision

8. ROI instances and illumination pattern retrieval

9. Fourier Ring Correlation (FRC) assessment of resolution

10. Precision and CRLB as a function of global phase

11. Precision and CRLB for 4 phase step dataset

12. Additional DNA-origami grid results

13. Impact of global phase on zero-background CRLB of the localization precision for a reduced phase scan range

14. SIMFLUX setup

15. Modulation estimation

16. High density (many non-spatially specific blinking fluorophores) initial pattern pitch estimation used as a starting point for low-density localization experiments

17. DNA origami structures

18. Timing Chart

19. Simulation study minimum filter

20. Data processing pipeline for pattern estimation and SIMFLUX localization

21. Deviation between SIMFLUX and SMLM localizations over the FOV

**Supplementary Figure 1.**
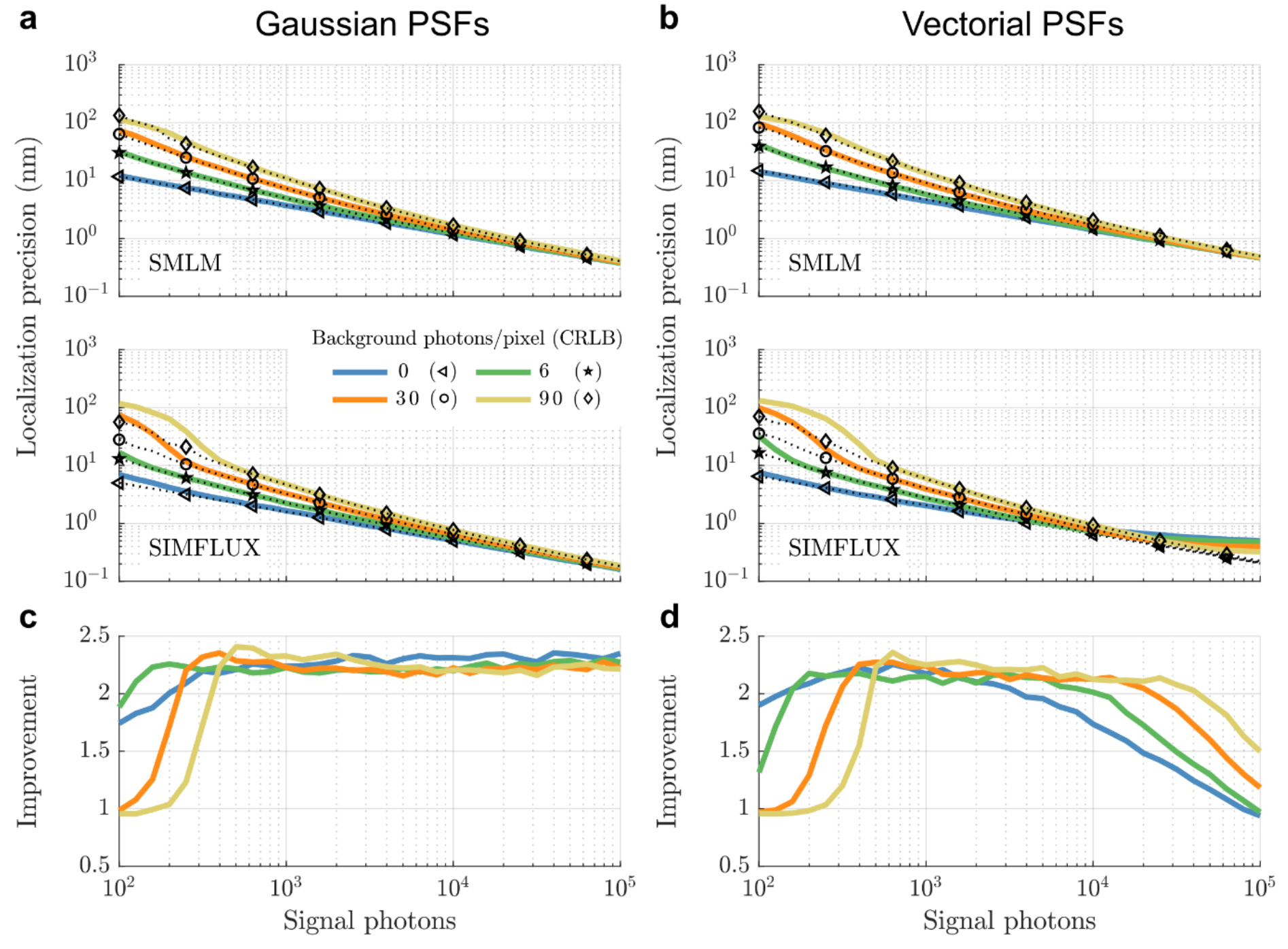
Impact of signal photon count on the localization precision of SIMFLUX in comparison to standard SMLM for different background levels. (**a-b**) Average lateral localization precision of SMLM and SIMFLUX fits using a Gaussian PSF model on ground truth data simulated with (a) Gaussian PSF (spot width *σ*_PSF_ = *λ*/4NA) and (b) vectorial PSF (see *Methods*), as a function of signal photon count for different background levels. The performance is at the CRLB except for SIMFLUX Gaussian PSF fitting on vectorial PSF ground truth simulations with very high signal-to-background-ratio (SBR), indicating a sensitivity to PSF model mismatch there. (**c-d**) Improvement factor of SIMFLUX localization precision over SMLM localization precision (= Δ*x*_SMLM_/Δ*x*_SIMFLUX_) for fitting with a Gaussian PSF model on ground truth data simulated with (c) Gaussian PSF and (d) vectorial PSF. Simulation parameters as described in *Methods*.

**Supplementary Figure 2.**
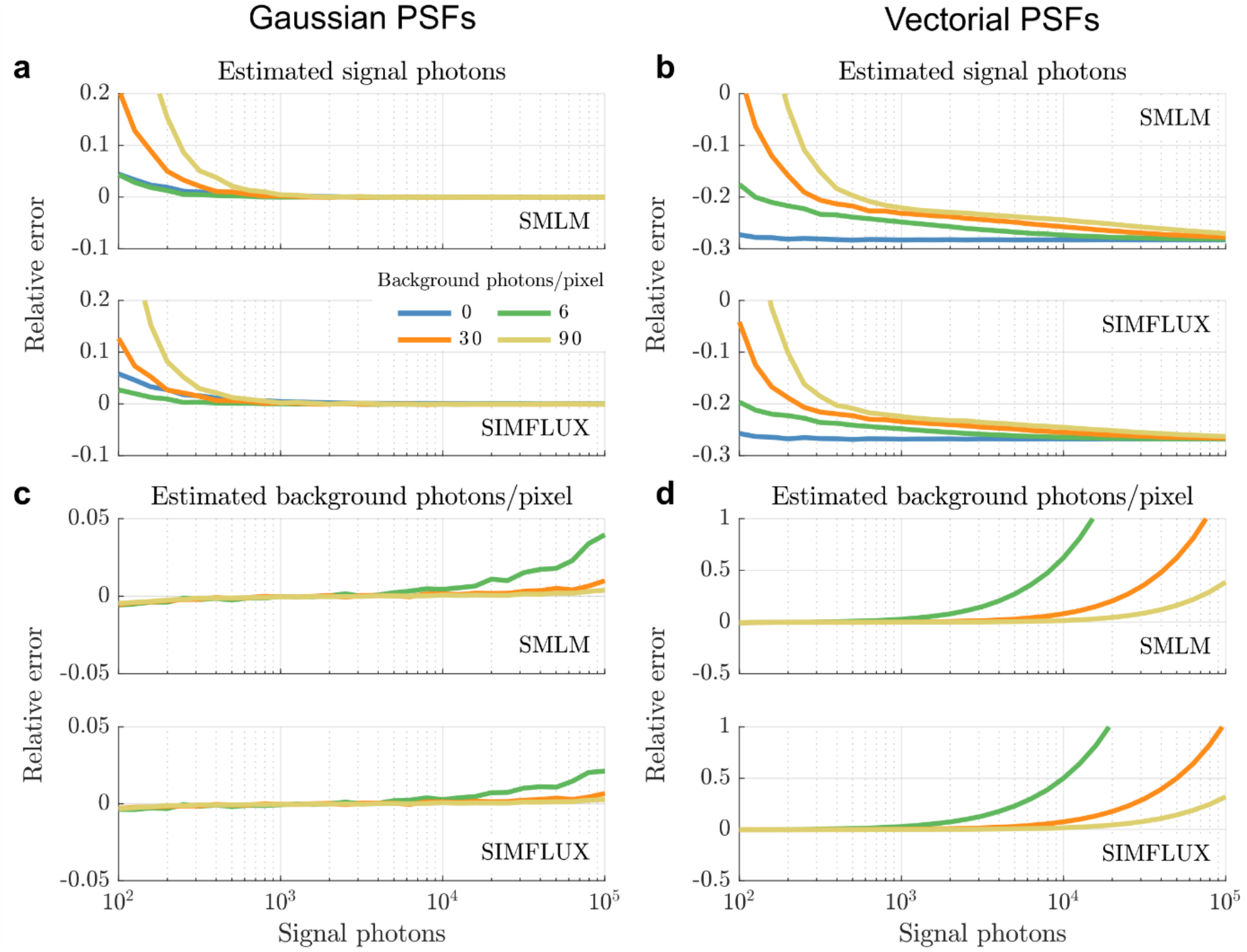
Impact of signal photon count on the photon count estimation of SIMFLUX in comparison to standard SMLM. (**a-b**) Relative error of the estimated signal photon count of SMLM and SIMFLUX fitting with a Gaussian PSF on ground truth simulated with (a) Gaussian (spot width *σ*_PSF_ = *λ*/4NA) and (b) vectorial PSFs (see *Methods*) as a function of signal photon count for different background levels. showing a perfect fit for Gaussian PSF fitting on Gaussian PSFs and a ~30% underestimation for a correct PSF model. (**c-d**) Similarly; relative error of the estimated background photons per pixel, indicating an overestimation for high signal-to-background ratio. The erroneous signal photon count estimation has an impact of the localization precision of SIMFLUX at high SBR, as shown in Supplementary Figure 1. Simulation parameters as in *Methods*.

**Supplementary Figure 3.**
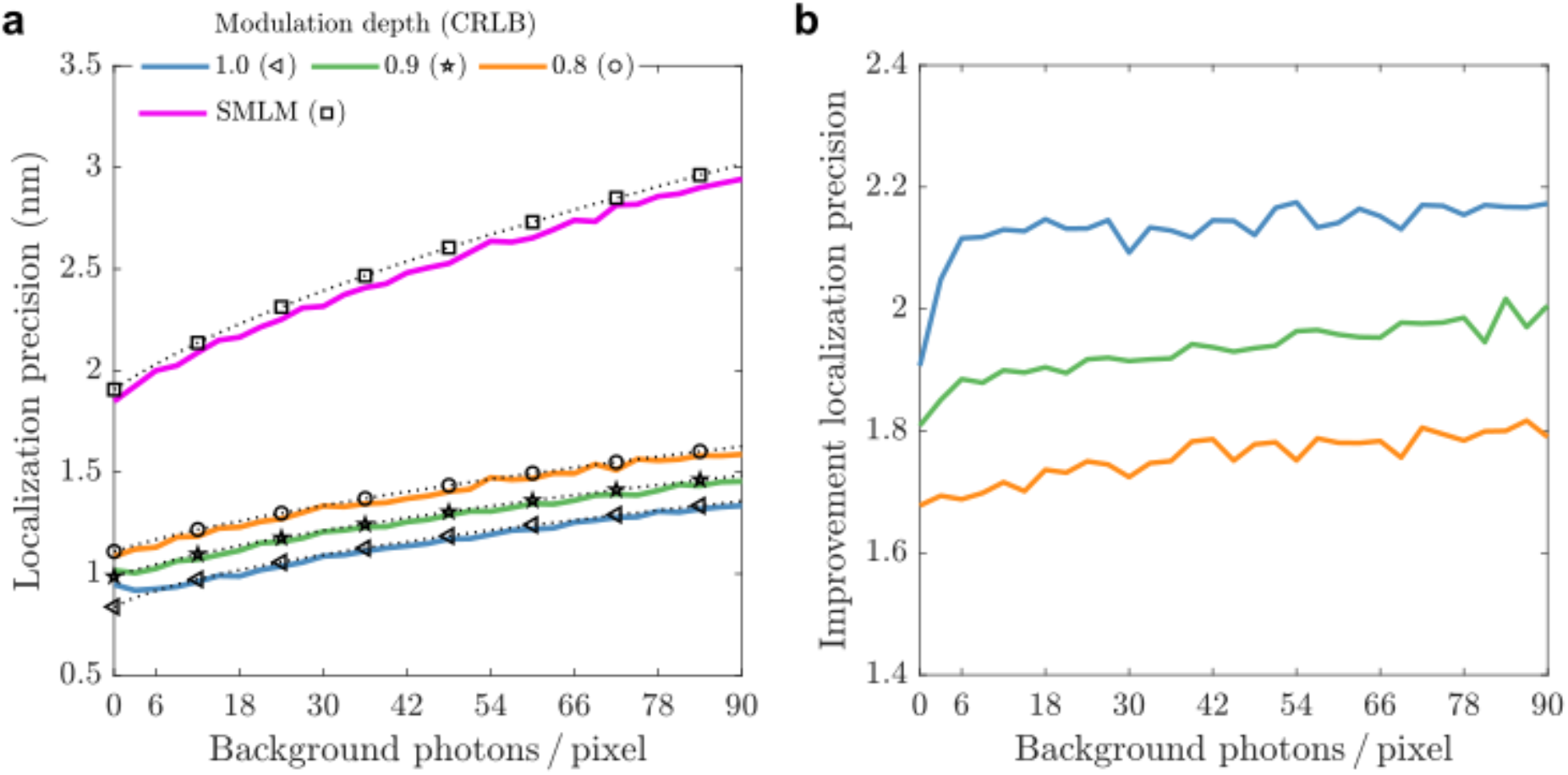
Impact of background photon count on localization precision of SIMFLUX in comparison to standard SMLM. **(a)** Average lateral localization precision of SMLM and SIMFLUX fitting with a Gaussian PSF model on vectorial PSF generated ground truth data as a function of background photons per pixel for three modulation depths. The performance is at the CRLB in the range of realistic background levels for the considered modulation depths. **(b)** Improvement of SIMFLUX over SMLM localization precision, showing a small impact of background on the improvement factor. The same relative impact of background on precision as in standard SMLM implies that SIMFLUX can be used under the same experimental conditions as SMLM.

**Supplementary Figure 4.**
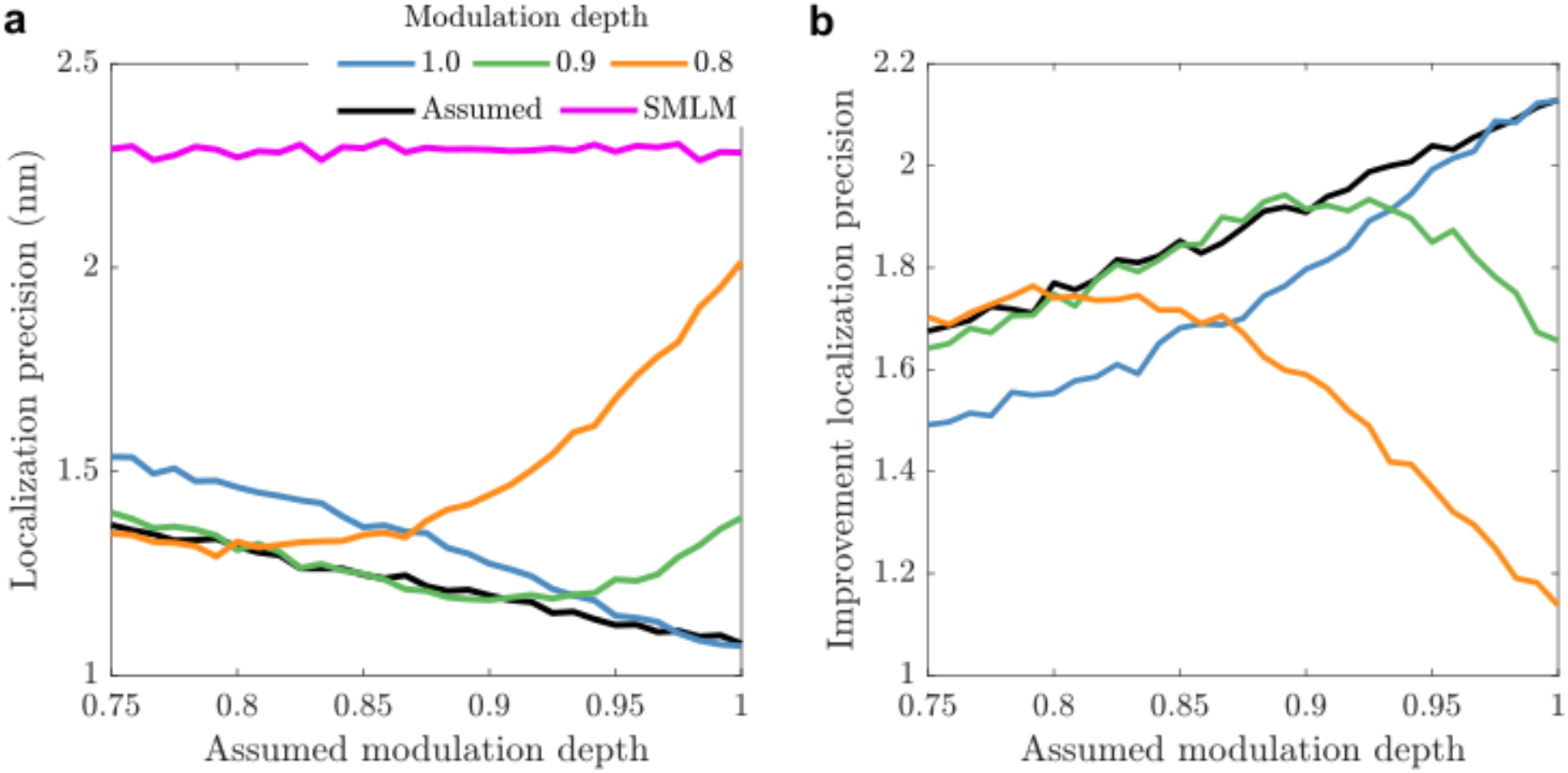
Assessment of illumination pattern modulation estimation on localization precision in SIMFLUX. **(a)** Average lateral localization precision in SIMFLUX as a function of an (incorrectly) assumed modulation depth in the SIMFLUX fit routine for different *actual* values of the modulation. For comparison the localization precision for SMLM on the summed simulation frames is plotted. Spots are simulated with 6000 signal photons and 30 background photons per pixel. **(b)** Improvement in SIMFLUX localization precision over SMLM localization precision as a function of an assumed modulation depth in the SIMFLUX fit routine for different actual values of the modulation, indicating that the modulation depth must be above about 0.9 for a ~2x improvement and must be known with a precision of about 0.05 for optimum fit results from comparing the green and black lines around 0.9.

**Supplementary Figure 5.**
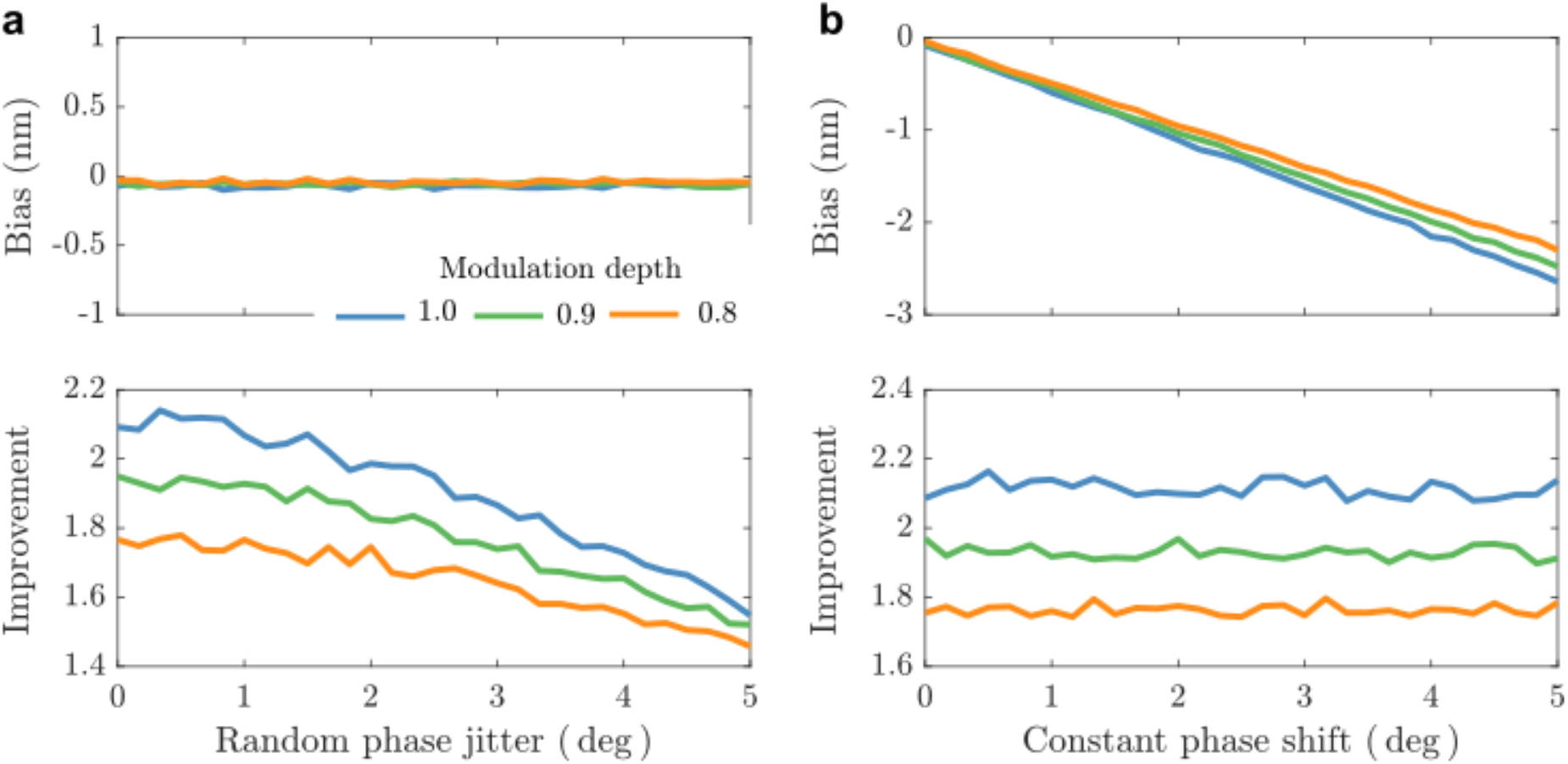
Impact of illumination pattern phase errors on localization precision and bias of SIMFLUX. **(a)** Average lateral localization bias of SIMFLUX and localization precision of SIMFLUX over standard SMLM for three modulation depths as a function of the standard deviation of random illumination pattern phase jitter, modelled by a Gaussian distribution. **(b)** Similarly; for a constant phase shift due to a potential bias in the phase estimation. The plots indicate a tolerable phase jitter of about 2 deg.

**Supplementary Figure 6.**
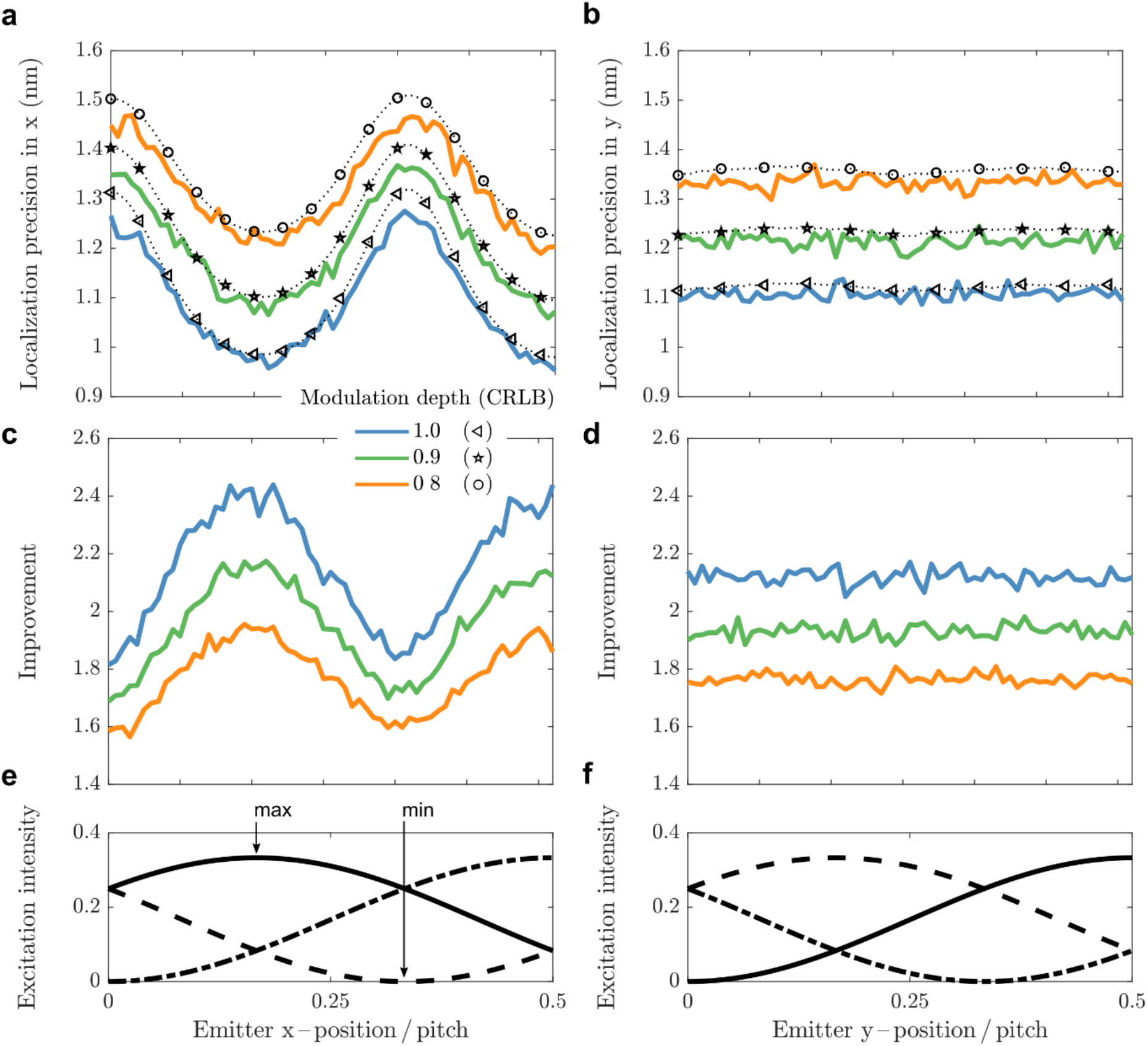
Impact of emitter position in the global phase pattern on localization precision in SIMFLUX. Simulated emitters with ground truth (**a**) *x*-positions fixed in the excitation pattern and (**b**) *y*-positions randomized in the excitation pattern over half the pitch. (**a-b**) SIMFLUX localization precision for different modulation depths as a function emitter position in the excitation pattern; the performance is at the CRLB. (**c-d**) Improvement of SIMFLUX over SMLM localization precision. (**e-f**) Normalized excitation patterns for perfect modulation in the (**e**) x-direction in which the x-coordinates are fixed and (**f**) y-direction where the y-coordinates are randomly distributed. The plots show the position dependent localization precision of SIMFLUX within the global phase pattern and the average localization precision. The best precision is achieved for an emitter position at the center of the illumination pattern maximum and worst for emitters located in a minimum. This is in line with theory (Supplementary Note and Supplementary Fig. 7), and deviates from MINFLUX because of the imperfect modulation depth.

**Supplementary Figure 7.**
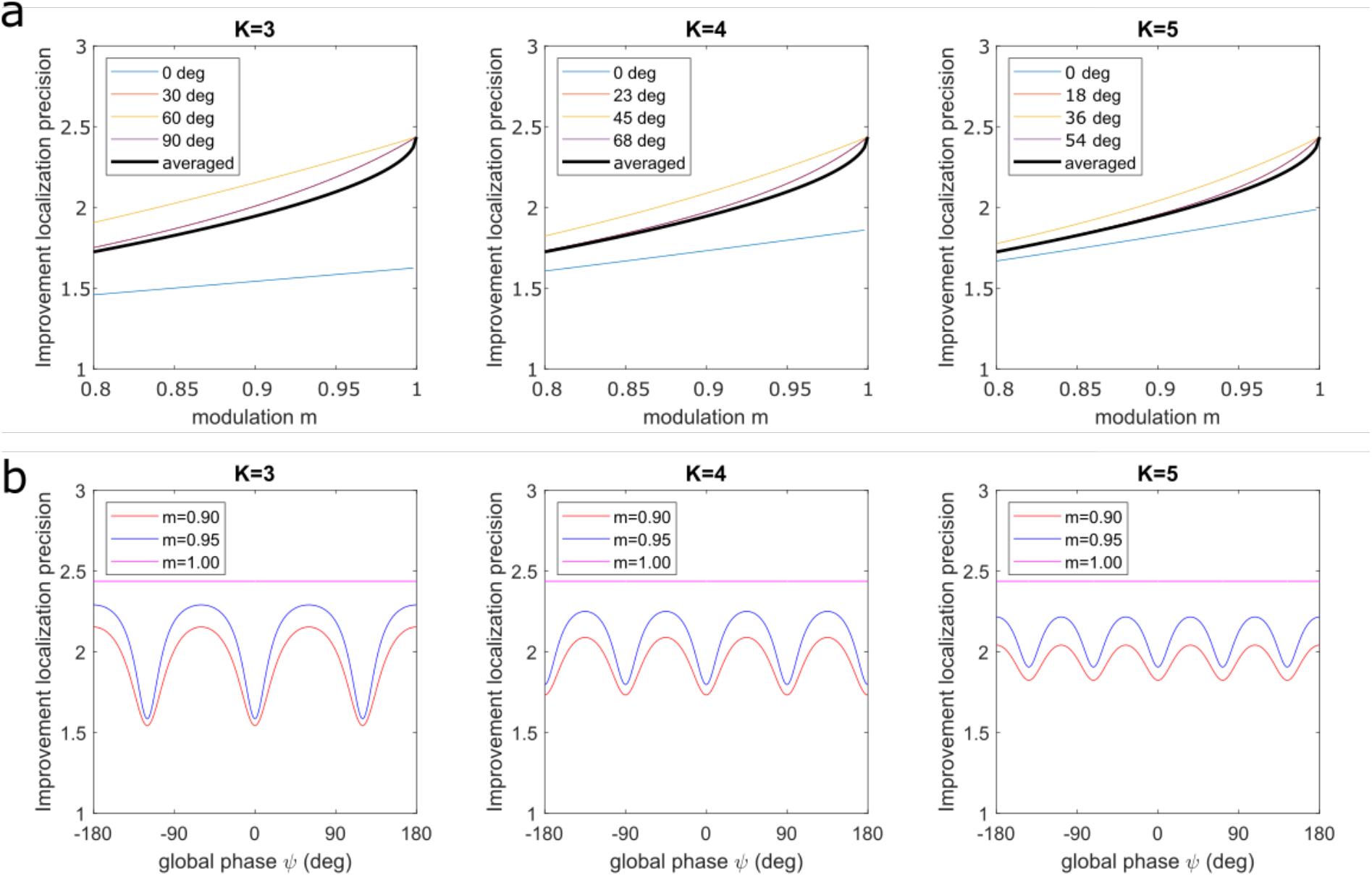
Impact of modulation depth and global phase on zero-background CRLB of the localization precision. **(a)** The improvement in localization precision of SIMFLUX over standard SMLM for different numbers of phase steps *K* and global phase of the molecule (measure for the position of the molecule from the intensity minimum of one of the patterns), as a function of modulation depth *m*, as well as the global phase averaged improvement factor. **(b)** The improvement in localization precision of SIMFLUX over standard SMLM for different numbers of phase steps *K* and modulation as a function of global phase. The plots indicate a steep dependence on *m* close to *m* = 1, in particular for the global phase where one of the images is acquired when the molecule is at the illumination pattern minimum. This agrees with the numerical simulations shown in Supplementary Fig. 6. The performance is worst for these cases in case the modulation is imperfect. For a larger number of phase steps, the variations in CRLB as a function of global phase strongly reduce, making the method more robust. In the computations we take a pitch to spot width ratio *p/σ* = 2.

**Supplementary Figure 8.**
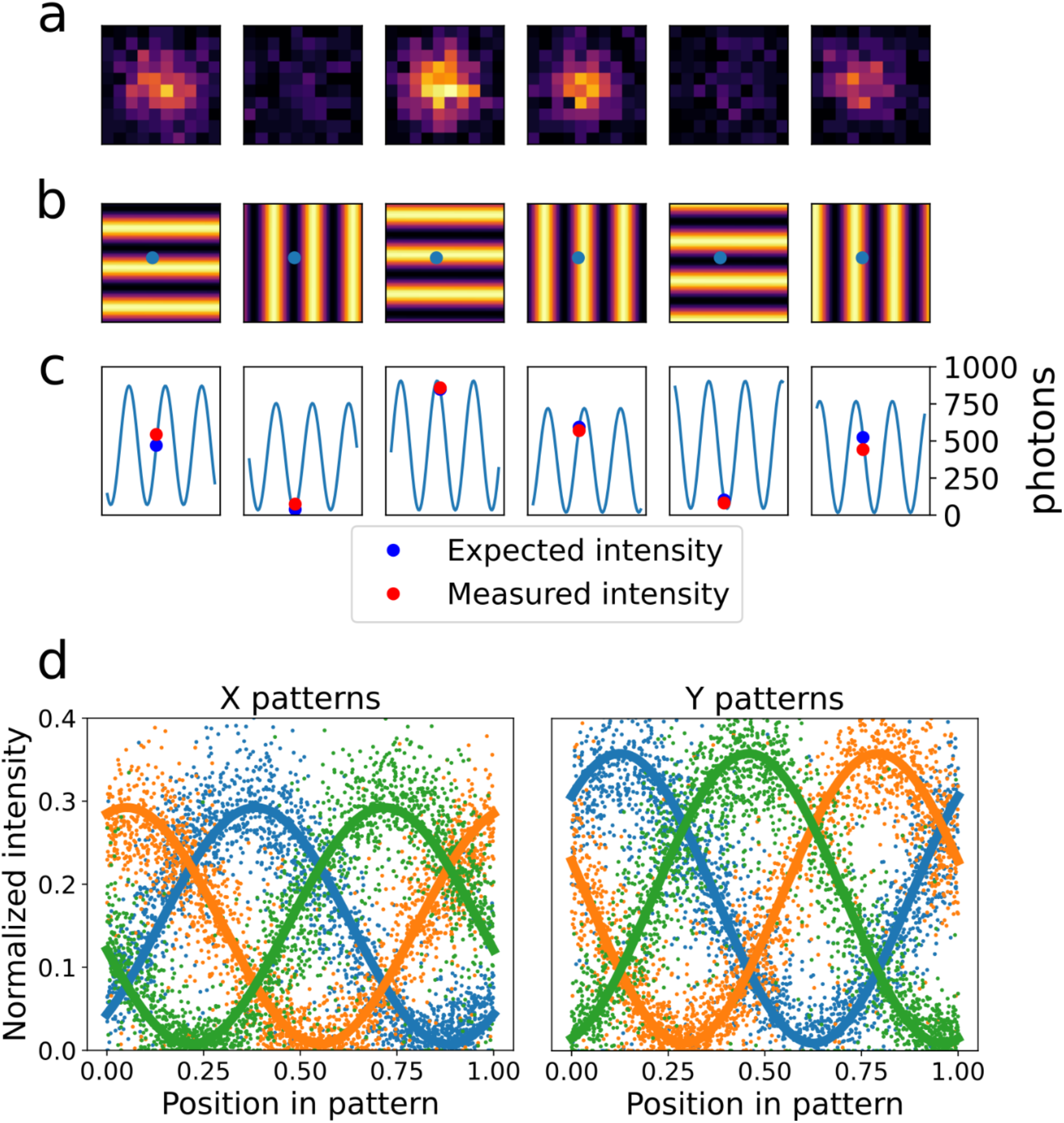
ROI instance and illumination pattern retrieval. **(a)** Single-molecule image (11×11 pixels of size 65 nm) over 6 subsequent frames, with estimated signal photon counts, showing the frame-to-frame variation in emission intensity caused by the shifting and rotating illumination pattern. **(b)** Retrieved illumination pattern with position of the molecule with respect to the pattern indicated. **(c)** Expected and actual signal photon count, showing a good match, within the margins of shot noise induced variations. **(d)** Fit of the sinusoidal illumination pattern through the entire set of localizations. The point clouds show the estimated ratio of signal photon count to total photon count (“normalized intensity”) over the *K* = 3 phase steps per pattern orientation as a function of *x* and *y*-position mapped into a single period of the illumination pattern. The pattern phases are estimated with a precision equal to 0.64, 0.59, 0.68, 0.24, 0.55, and 0.41 deg (order pertaining to patterns as shown in second row), assessed with the procedure outlined in Methods.

**Supplementary Figure 9.**
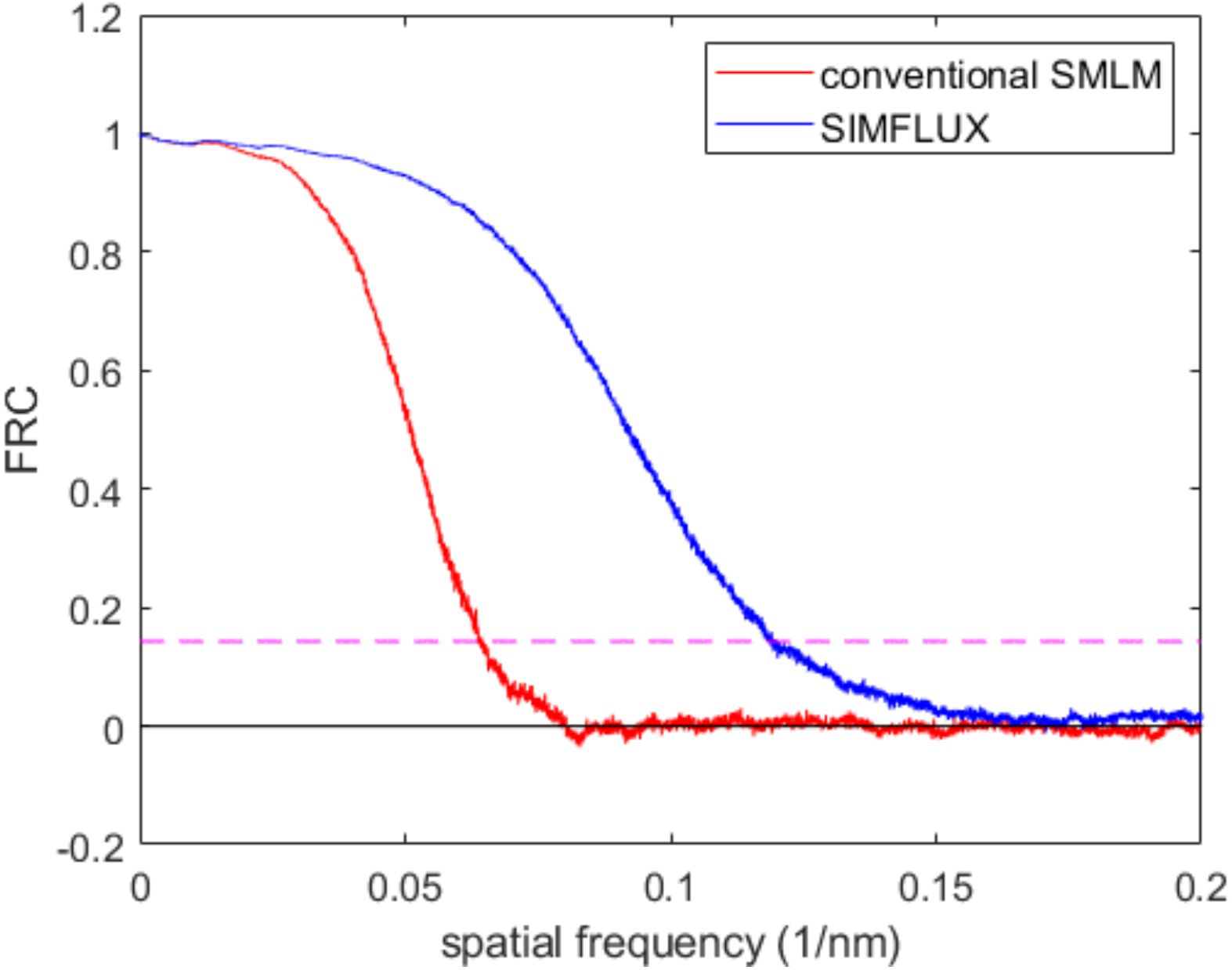
Fourier Ring Correlation (FRC) assessment of resolution. The FRC-curve for SIMFLUX shows a clear improvement in resolution over conventional SMLM. The resolution values, as determined from the intersection with the 1/7-threshold (dashed line), are 8.4 nm (SIMFLUX) and 15.5 nm (SMLM), with a typical uncertainty better than 0.5 nm, implying an improvement with a factor 1.8. The two image halves are found by randomly selecting localizations to the two subsets. This gives rise to FRC curves largely determined by the localization precision, eliminating correlations arising from having multiple localizations from the same binding site (“spurious correlations”) would result in an FRC-curve determined by the structure of the sparsely distributed binding sites^19^. The split datasets are used to generate reconstructions on a 2 nm super-resolution pixel grid (super-resolution pixel size must be less than about 0.25× the FRC-resolution) by the histogram binning method.

**Supplementary Figure 10.**
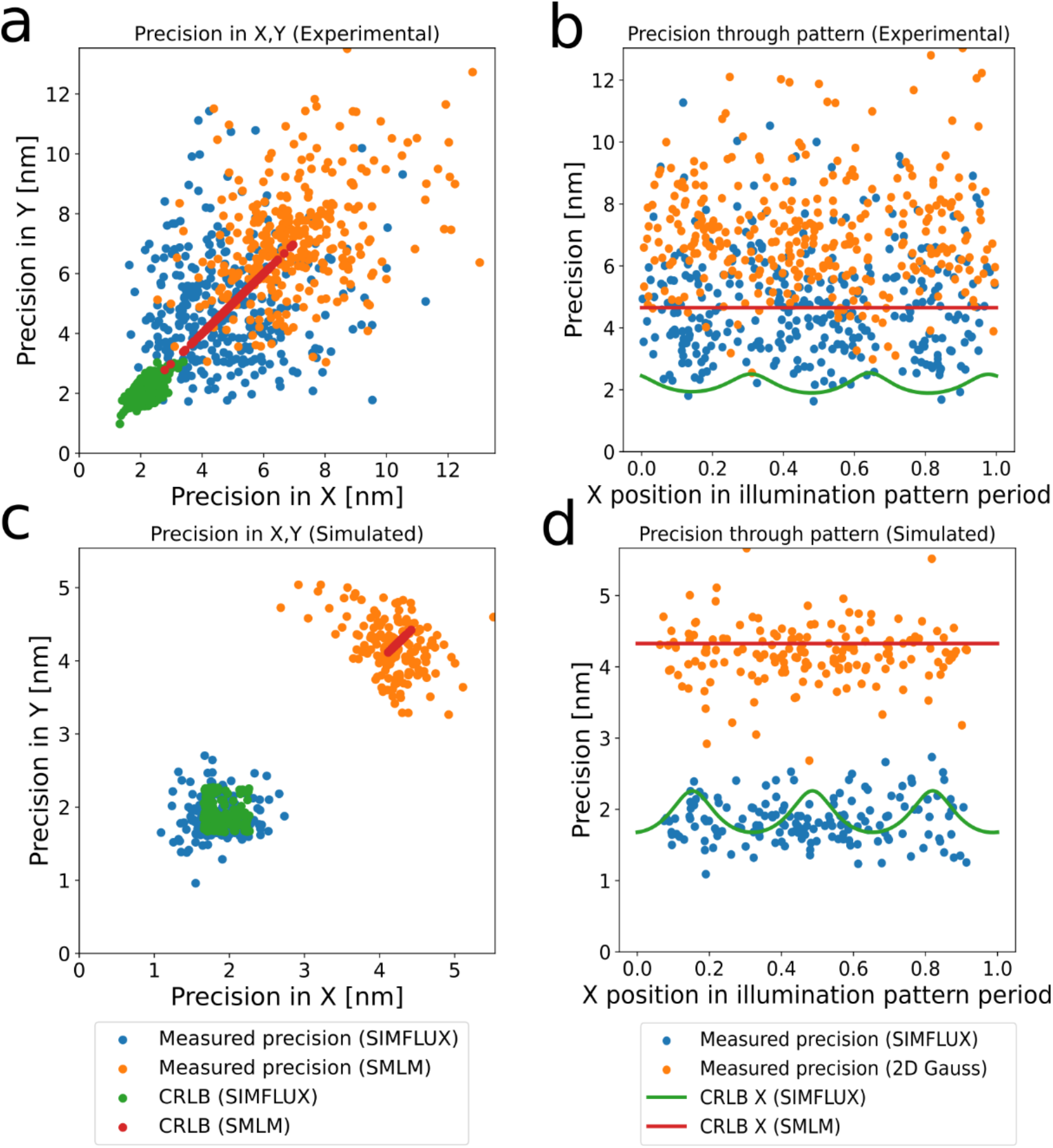
Precision and CRLB as a function of global phase. **(a)** Measured precision in *x* and *y* for 431 analysed clusters of localizations for the 80 nm nanoruler dataset of Fig. 2, in comparison to the average CRLB over the localizations within each cluster. The clusters correspond to the binding sites of the nanorulers, the precision is quantified by the standard deviation of the set of localizations within each cluster. The total set of localizations has a median emitter intensity of 1294 photons, cumulative over the 6 frames, and a background of 8.4 photons/pixel, cumulative over the 6 frames. **(b)** Measured precision and CRLB as a function of the *x*-coordinate, where the *x*-values are mapped into a single period of the illumination pattern (the “global phase”). The median of the distribution of precision values is 4.5 nm for SIMFLUX and 6.9 nm for conventional SMLM, indicating an improvement factor of 1.5. The performance is not on par with the CRLB, probably due to a residual drift of around 4 nm over the full dataset. **(c,d)** Same as (a,b), for a simulated full-FOV dataset (signal photon count 1406, background/pixel 7.6), indicating a precision improvement factor of 2.3, and a performance on par with the CRLB.

**Supplementary Figure 11.**
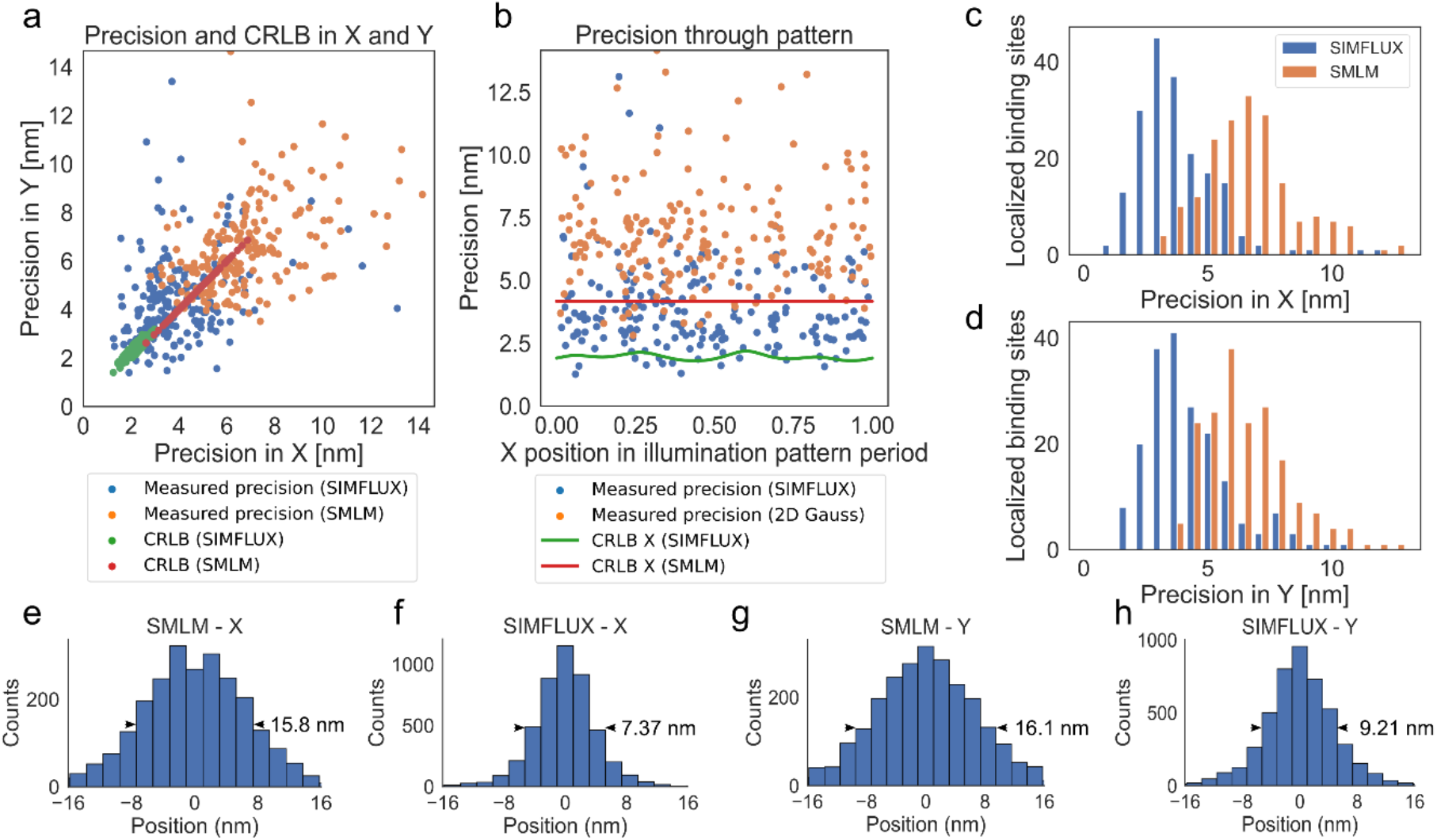
Precision and CRLB for SIMFLUX on 80 nm nanorulers with *K* = 4 phase steps. **(a)** Measured precision in *x* and *y* for 194 analysed clusters of localizations, in comparison to the average CRLB over the localizations within each cluster. **(b)** Measured precision and CRLB as a function of the *x*-coordinate, where the *x*-values are mapped into a single period of the illumination pattern (the “global phase”). The total set of localizations has a median emitter intensity of 1645 photons, cumulative over the 8 frames, and a background of 11.3 photons/pixel, cumulative over the 8 frames. **(c,d)** Distribution of precision values, with median 3.7 nm for SIMFLUX and 6.4 nm for conventional SMLM (average over *x* and *y*), indicating an improvement factor of 1.7. **(e-h)** Histograms of the total set of localizations, with average FWHM equal to 16.0 nm (SMLM) and 7.8 nm (SIMFLUX), indicating an improvement factor of 2.0. Comparison to the similar plots for *K* = 3 phase steps in Supplementary Fig. 9 indicates a comparable precision and a somewhat larger improvement factor.

**Supplementary Figure 12.**
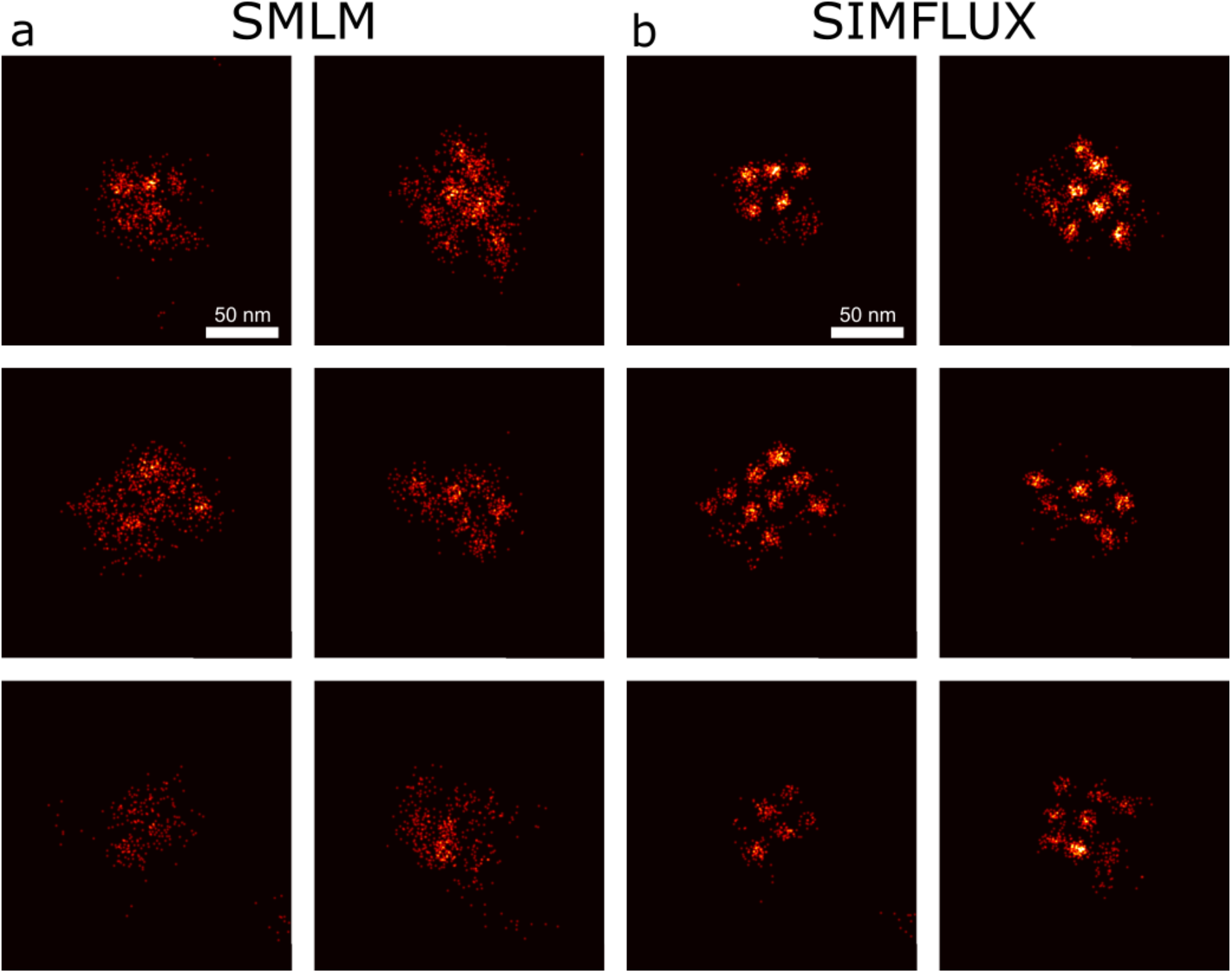
Additional DNA-origami grid results. **(a)** Conventional SMLM and **(b)** SIMFLUX images of selected instances of the 20 nm DNA-origami grid structures. Imperfect labelling prevents all 4×3 binding sites to be visible. The grid structure is not resolved in SMLM, but clearly distinguishable in SIMFLUX.

**Supplementary Figure 13.**
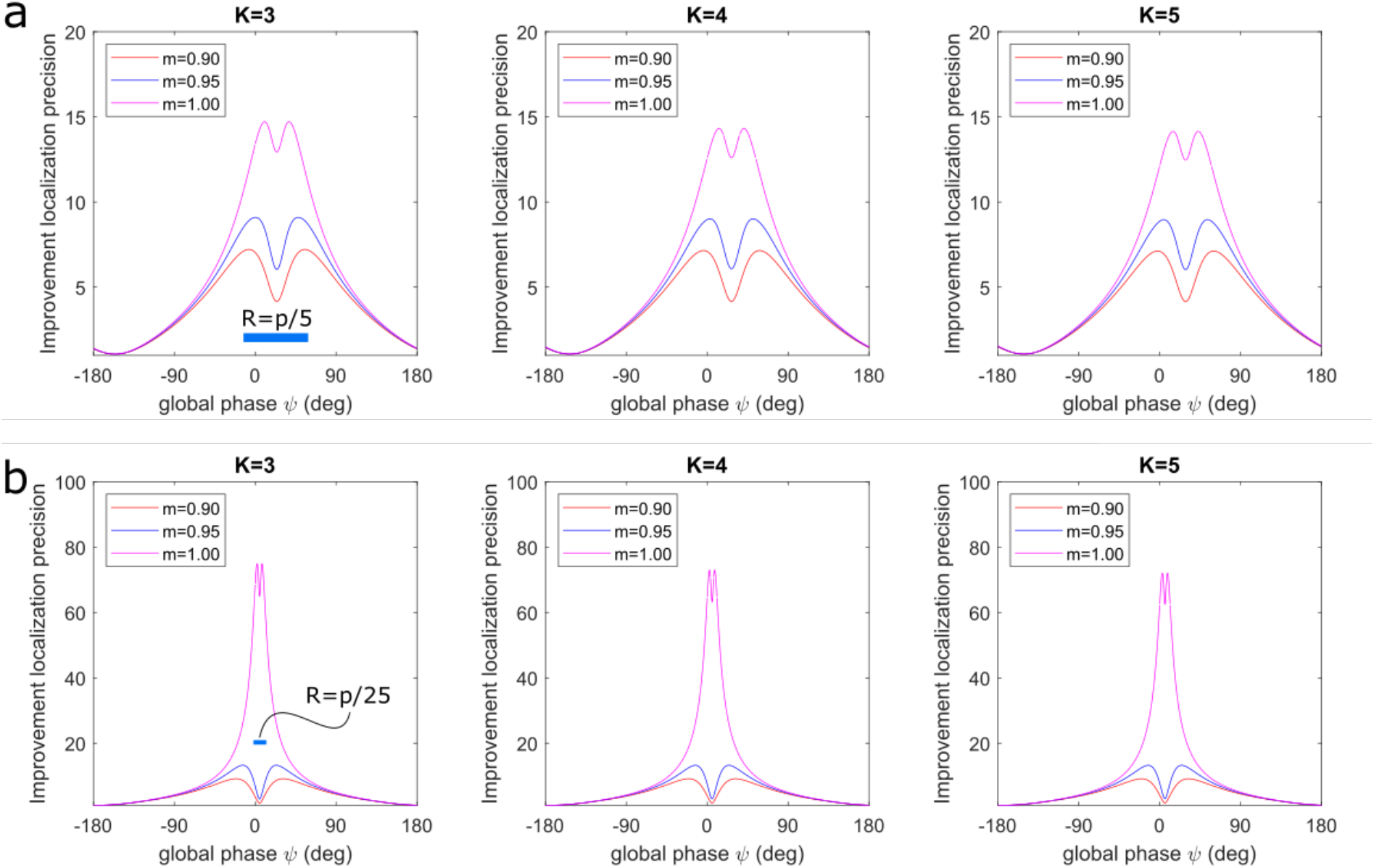
Impact of global phase on zero-background CRLB of the localization precision for a reduced phase scan range. **(a)** Improvement in localization precision of SIMFLUX over standard SMLM as a function of global phase for three different modulation values for a scan range *R* = *p*/5, with *p* the pattern pitch. **(b)** Same for a scan range *R* = *p*/25. In the computations we take a pitch to spot width ratio *p*/*σ* = 2. The plots show a huge improvement factor when the molecule is placed within the range of positions where the illumination pattern minimum is scanned. Interestingly, for an imperfect modulation, the improvement factor reaches a maximum around 10, for a position of the molecule at the edge or just outside the range of positions where the illumination pattern minimum is scanned. Close to the illumination pattern maximum the improvement factor then collapses to a value close to one. This is in line with the performance for the full scan range *R* = *p*, shown in Supplementary Figs. 6 and 7.

**Supplementary Figure 14.**
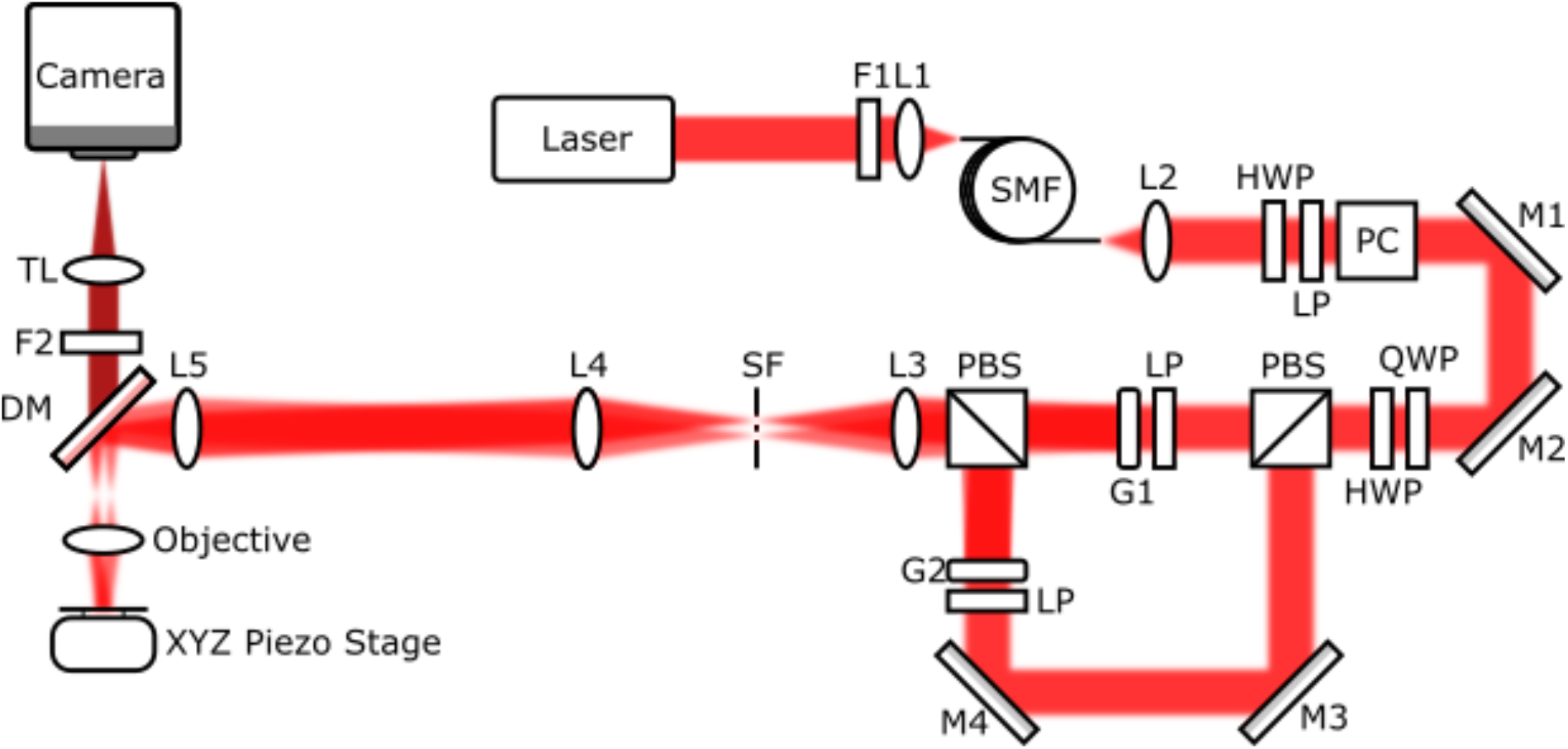
SIMFLUX setup. Laser 640 nm, F1 excitation filter, L1 fiber coupling lens, SMF polarization maintaining single mode fiber, L2 fiber collimation lens, HWP zero order half wave plate 633 nm, QWP zero order quarter waveplate 633 nm, LP glan-laser linear polarizer, PC Pockels cell, M1-4 aluminium steering mirrors, PBS polarizing beam splitter, G1,2 binary phase gratings mounted on piezo stages, L3 75 mm relay lens, SF spatial filter, L4 350 mm relay lens, L5 180 mm relay lens, Objective 1.49 NA TIRF, XYZ Piezo Stage 100×100×100 nm travel range piezo stage, DM dichroic long pass mirror, F2 emission filter, TL tube lens, Camera sCMOS Hamamatsu Orca Flash 4.0 V2.

**Supplementary Figure 15.**
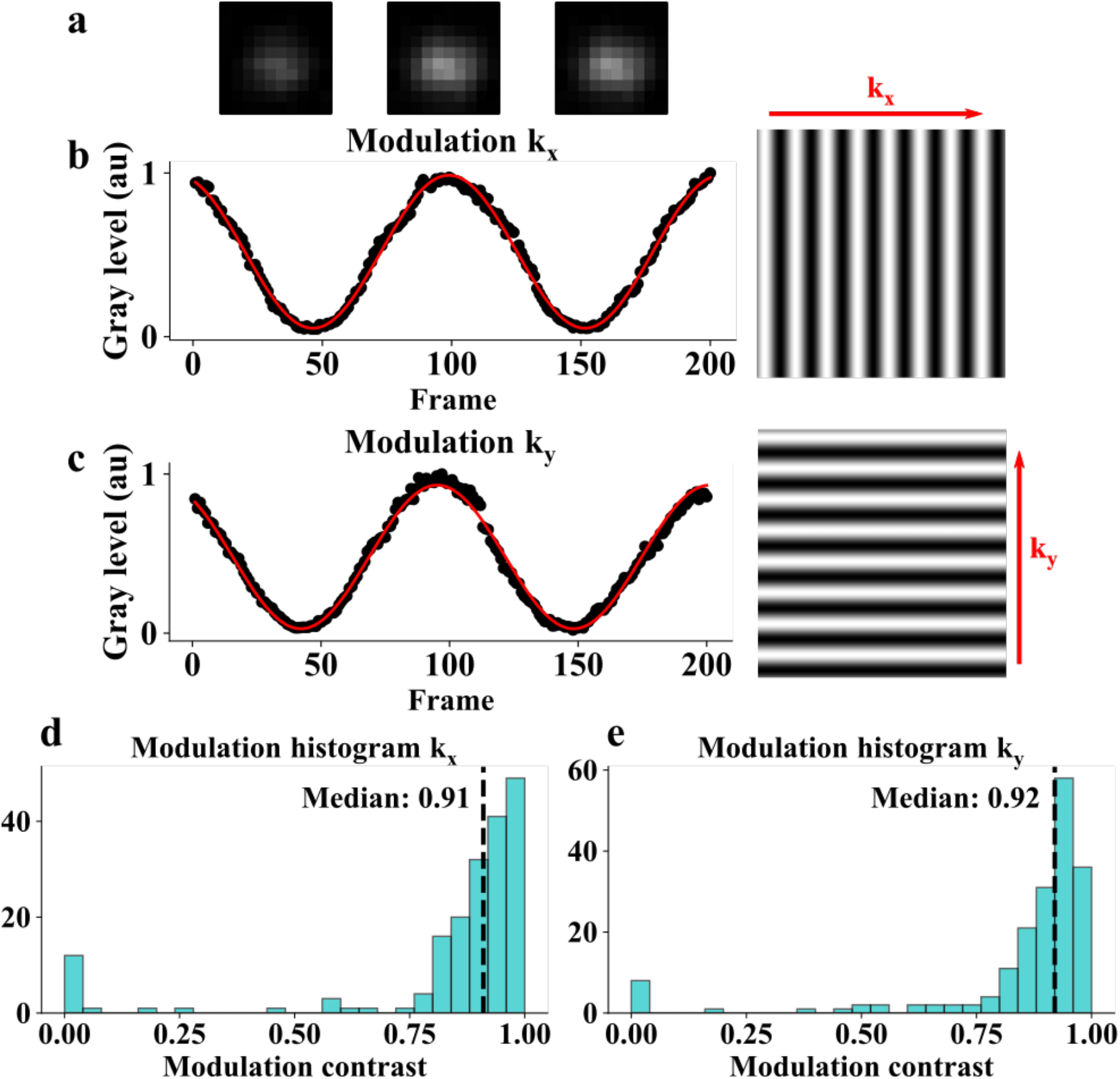
Modulation estimation. **(a)** Cropped image of a bead in a 10×10 pixel ROI at three phases that roughly correspond to the frame number on the *x* axis of the *x* modulation plot **(b,c)** Measured brightness of a 20 nm bead as a function of camera frame where for each frame the phase grating is shifted by 40 nm, out of a 8.496 μm period, in the grating plane with the piezos. ROIs of size 11×11 pixels were automatically segmented from a 26 μm field of view and their summed ADU (minus the background) was fit with a sinusoidal curve as described in the Methods section with an *R*^2^ > 0.98 for both directions. **(d,e)** Estimated modulation from 184 ROIs over the 26 μm field of view.

**Supplementary Figure 16.**
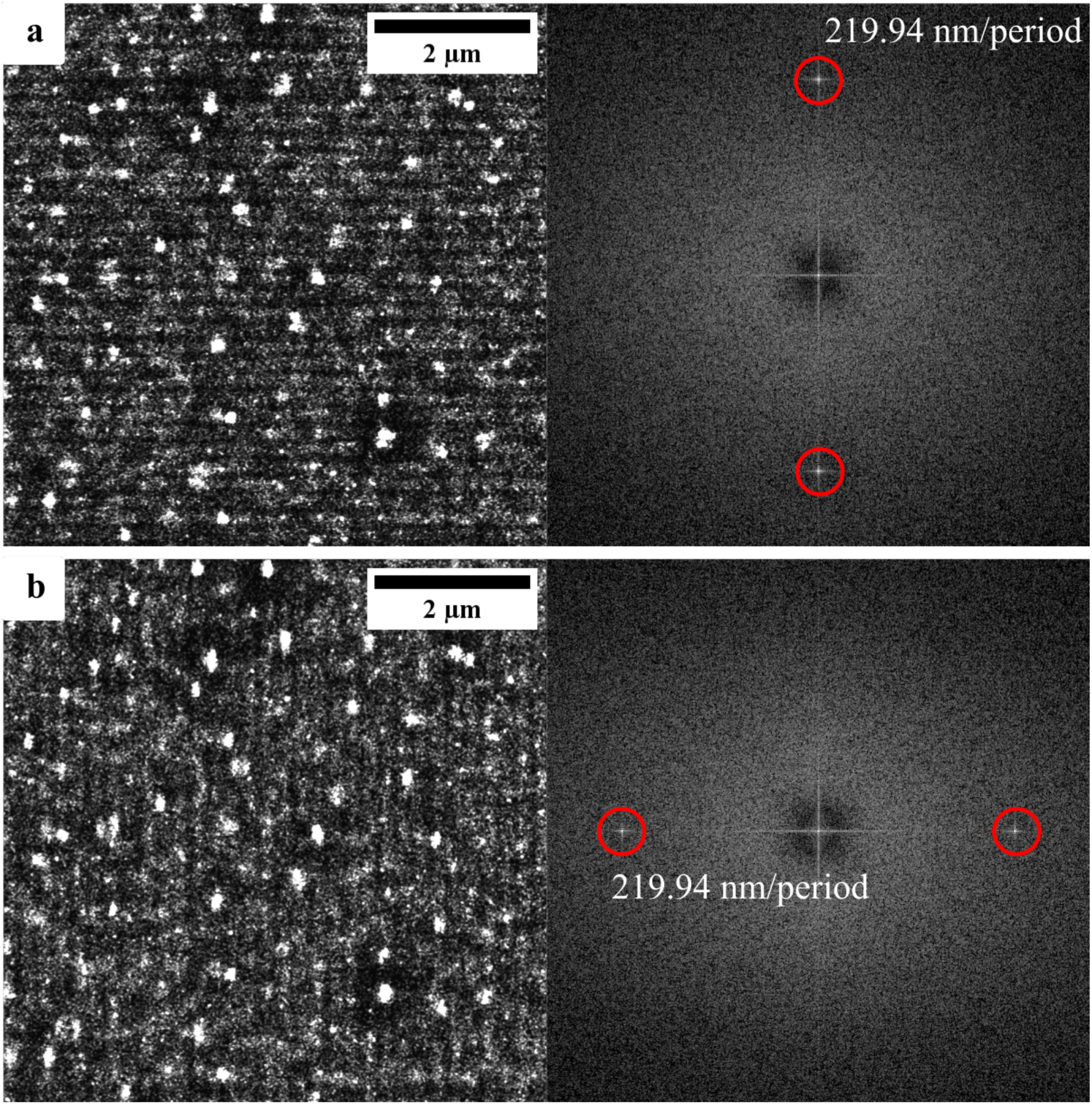
High density (many non-spatially specific blinking fluorophores) initial pattern pitch estimation used as a starting point for low-density localization experiments, **(a)** 600×600 pixel cropped region of interest from localized high-density fluorophores illuminated with a periodic pattern, and Fourier transform of the image for the *x*-oriented pattern. **(b)** Same for the *y*-oriented pattern. The estimated pitch is displayed in the image, and agrees well with the expected value 219.9 nm (see Methods).

**Supplementary Figure 17.**
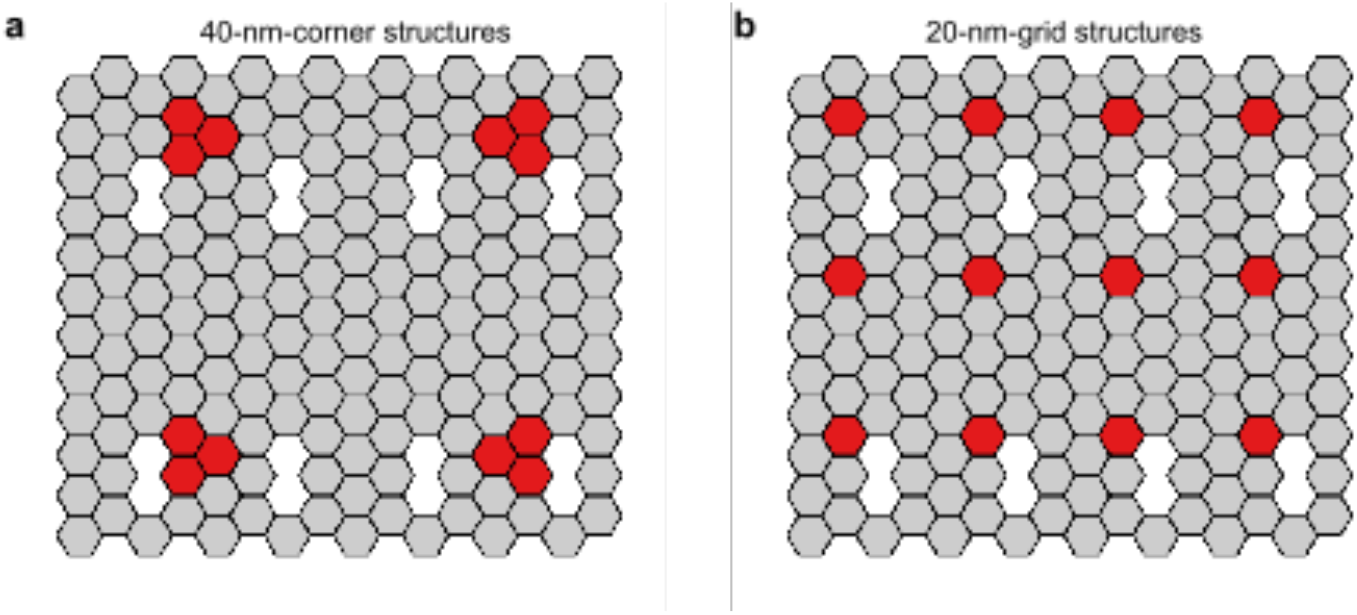
DNA origami structures. **(a)** 2×2 square with 40 nm spacing between binding sites. **(b)** 4×3 grid with 20 nm spacing. Not all strands/binding sites are present in the image, however, due to a limited incorporation efficiency.

**Supplementary Figure 18.**
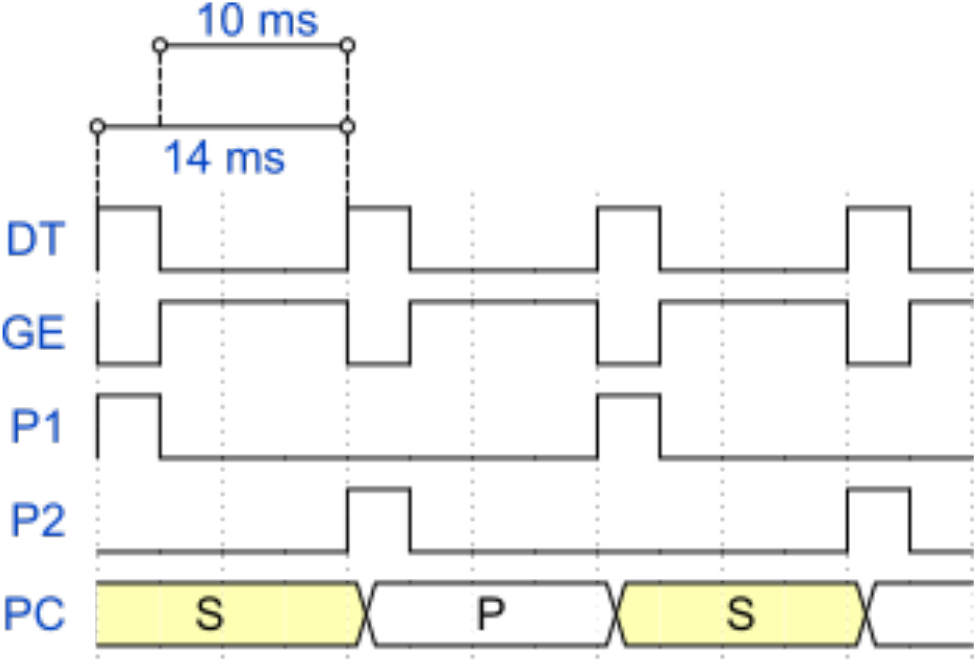
Timing chart. DT digital trigger, GE global exposure, P12 piezo1,2, PC Pockels cell. The digital trigger (DT) causes a global exposure (GE) event to occur on the camera. The timings between the DT trigger (10 ms and 14 ms can be adjusted). GE is high when all pixels experience the same amount of exposure. Only acquiring images and illuminating the sample during the global exposure ensures that there is an even flux of photons across the field of view and that there is no cross-talk between our two imaging arms. The piezoelectric stages P1 and P2 are set to translate a by a user defined amount every other DT high trigger in between global exposure events, ensuring that the piezo gratings are not moving during their acquisition period. Again, the Pockels cell (PC) switches between the S and P beam paths by inducing a half wave voltage every DT high trigger event.

**Supplementary Figure 19.**
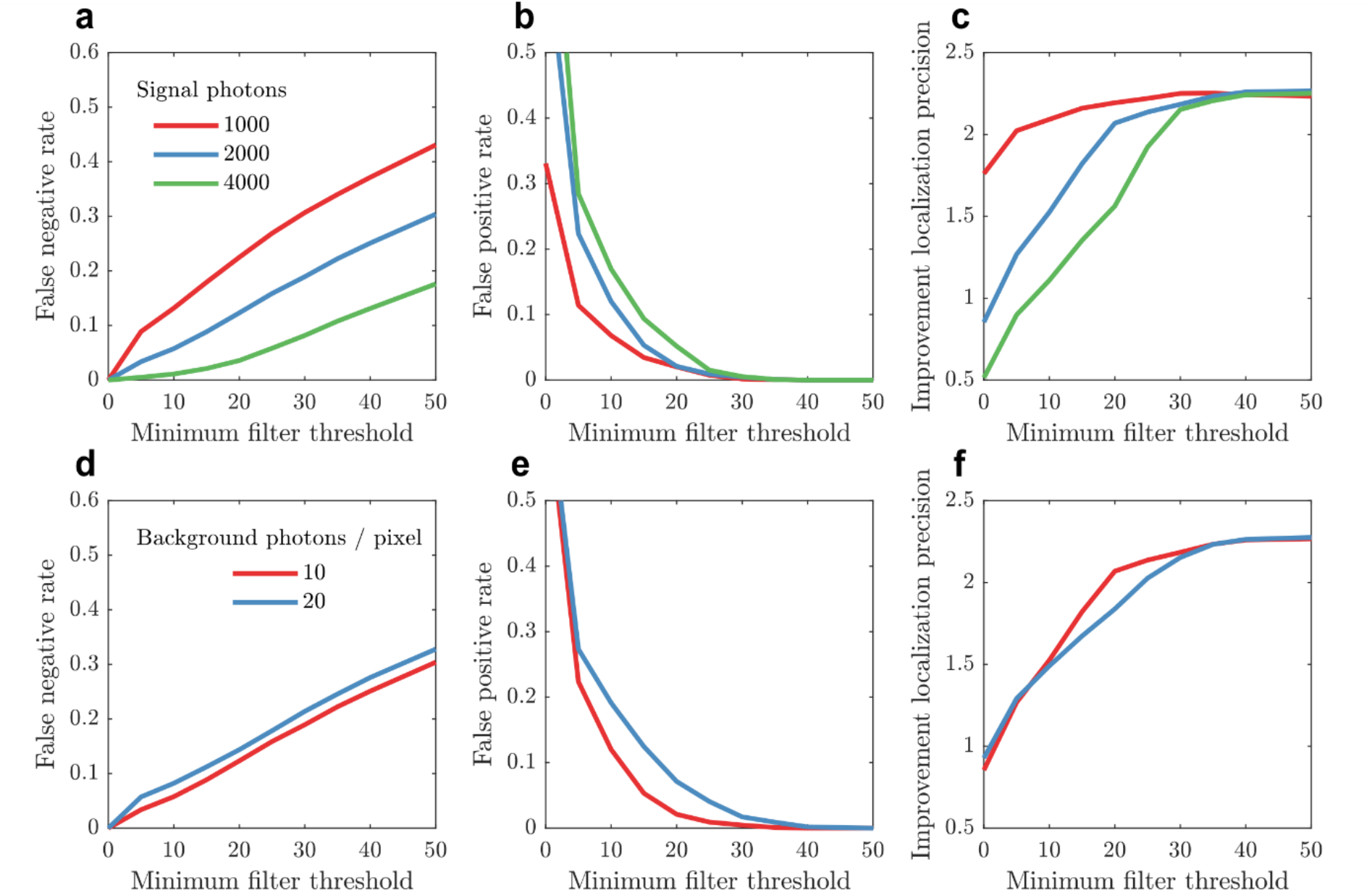
Simulation study of using a minimum filter for on/off transition estimation. **(a)** Ratio of molecular on-events that are on for all 6 phase step frames but incorrectly rejected by the minimum filter to the total number of on-events that last the full 6 frames (false negative rate) as a function of the minimum filter threshold for different signal photon counts. The cumulative background over the 6 frames is 10 photons/pixel. **(b)** Ratio of molecular on-events with an on/off transition after the first or before the last of the 6 frames and that are incorrectly accepted to the total number of on-events that do not last the full 6 frames (false positive rate) as a function of the minimum filter threshold for different signal photon counts. **(c)** Improvement factor of SIMFLUX localization precision compared to the SMLM localization precision computed over all spots in the simulated FOV as a function of the minimum filter threshold for different signal photon counts. **(d)** False negative rate as a function of minimum filter threshold for different background levels. **(e)** False positive rate as a function of minimum filter threshold for different background levels. **(f)** Precision improvement factor as a function of minimum filter threshold for different background levels. The cumulative signal photon count over the 6 frames is 2000. Simulation parameters as described in *Methods*.

**Supplementary Figure 20.**
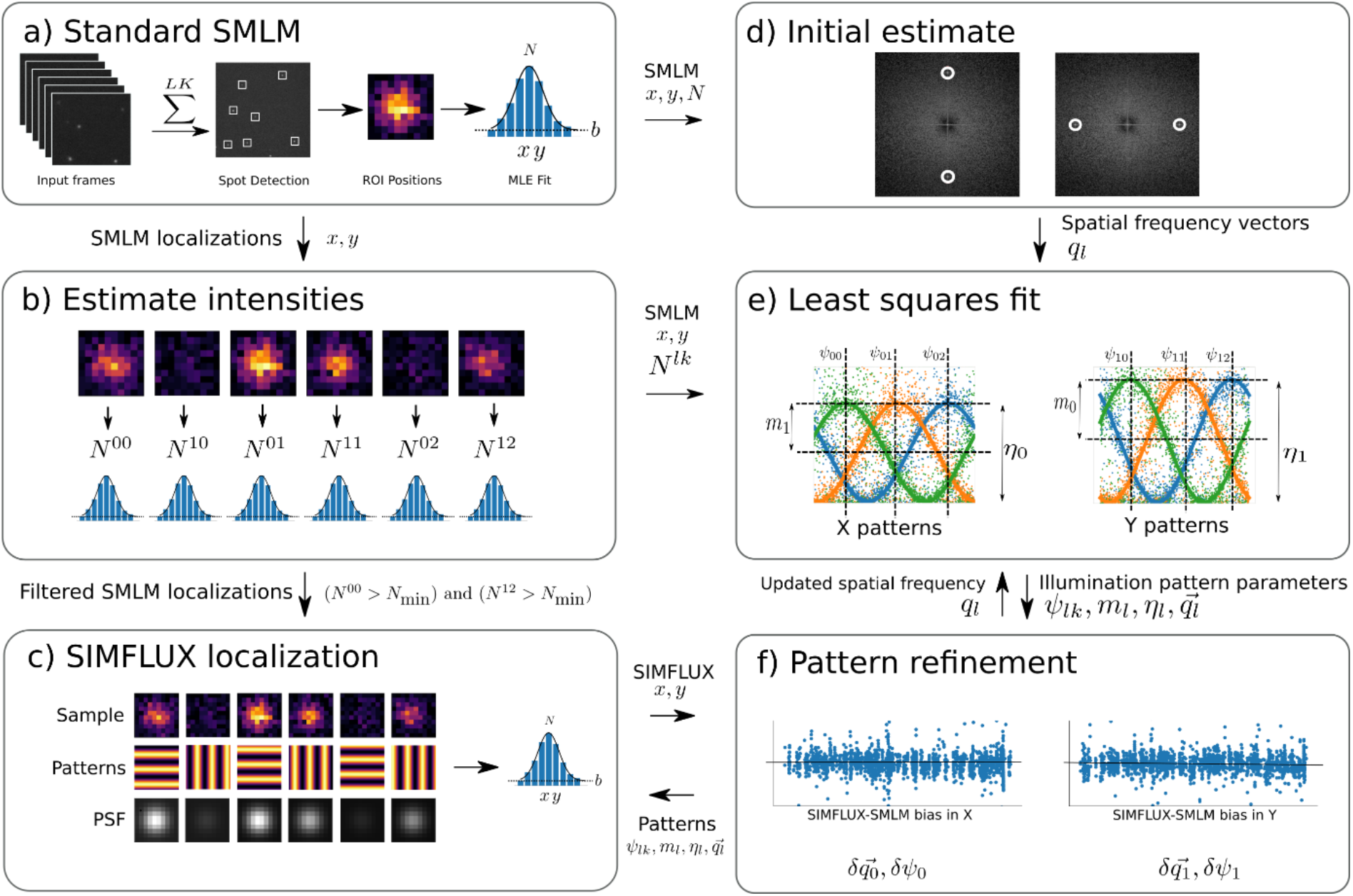
Data processing pipeline for pattern estimation and SIMFLUX localization. **(a)** The raw frames from the camera are converted to photon counts, and summed in blocks of 6 frames. The conventional localization microscopy pipeline is then used to perform spot detection and 2D Gaussian localization. **(b)** Filtering of the localizations. The signal intensity (photon count) on the first and the last frame of an on-event is estimated using a 2D Gaussian fit, during which the molecule *x, y*-position is kept fixed. Molecules for which the signal photon count on the first or last frame is lower than a predetermined threshold *N_min_* are rejected. **(c)** SIMFLUX localization. The 6 ROI frames are fitted to a model that uses both the illumination pattern information and the 2D spot center, resulting in a higher precision. **(d)**. Initial estimates for the illumination pattern pitch and angles are found by locating peaks in the Fourier domain of the rendered standard SMLM image. The SMLM localizations are rendered into a super resolution image with a zoom factor of 6 compared to the camera pixel size. A 2D Fourier transform is then used with zero padding to zoom in on the peak. The peak is then fitted with a 1D quadratic fit in both *q_x_* and *q_y_*-directions to calculate to subpixel peak position. **(e)** Using the estimated spatial frequency vectors 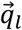, intensities 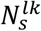 and SMLM localizations (*x_s_, y_s_*), the phases *ψ_lk_*, modulation depths *m_l_* and relative intensities *η_l_* of the *x* and *y*-pattern are computed using a least square fit as described in the methods section. **(f)** The pitch and phase offset is refined by doing a least squares line fit on the difference between the SMLM and SIMFLUX localizations in both modulation orientations. A nonzero slope of this line will correspond to an error in the pitch, and a nonzero offset will correspond to a bias shared between all phases of the modulation orientation.

**Supplementary Figure 21.**
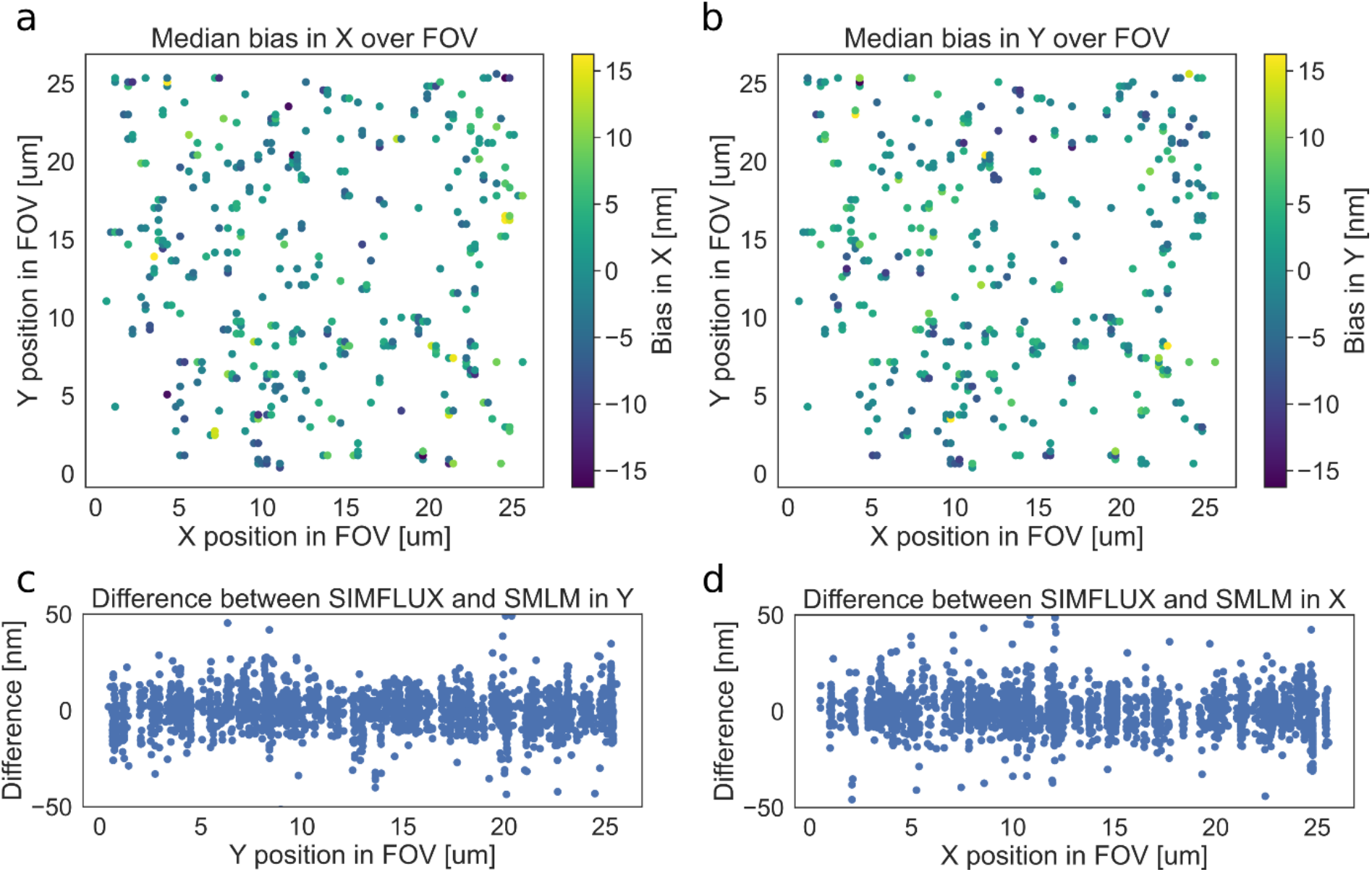
Deviation between SIMFLUX and SMLM localizations over the FOV. **(a,b)** Difference between the SIMFLUX and conventional SMLM position estimate for 377 analysed clusters of localizations, projected on the *x*-oriented and *y*-oriented pattern directions. The data is averaged over all localizations within each cluster. **(c,d)** show the cross-sections of (a,b). The plots show no systematic error over the FOV, indicating that the iterative procedure of estimating the pitches and orientations of the illumination patterns converges to a uniform description of the sinusoidal illumination pattern. The rms value of the SIMFLUX-SMLM bias over the full dataset is 12.9 nm (*x*) and 13.1 nm (*y*), on the order of the localization uncertainty.

## Supplementary Note

### 1. Image formation model

We have a sequence of *k* = 1,2,…, *K* illuminations for orientations *l* = 1,2,…, *L* with a harmonic intensity profile *P*(*φ*) as a function of pattern phase *φ* that is displaced according to phase offsets *ψ_lk_* such that:

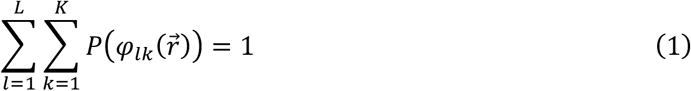

with the phase:

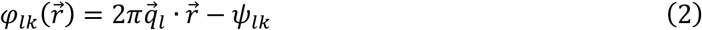

Here the spatial frequency vectors are:

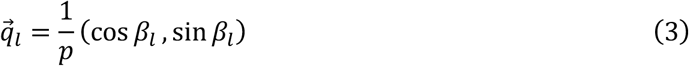

where *β_l_* = *πl*/*L* + *β*_0_, with *β*_0_ a global angular offset. The Point Spread Function (PSF) is 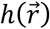, and is assumed to be a Gaussian:

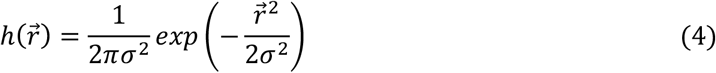

with *σ* the spot width. The expected photon count on pixel *j* is:

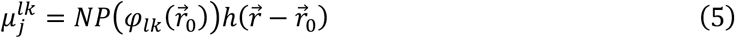

with 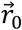 the emitter position and *N* the total photon count. We will consider a sinusoidal illumination pattern:

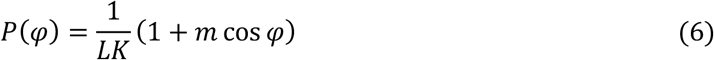

with *m* the modulation. For a perfect modulation *m* = 1, we find:

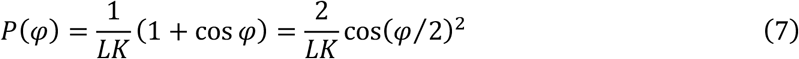

The phases of the different illumination patterns are assumed to be equidistant. For a full 2*π* phase scan this implies:

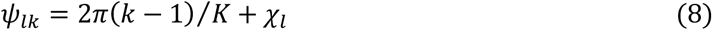

where *χ_l_* is the phase offset of the patterns in direction 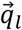. We then find that:

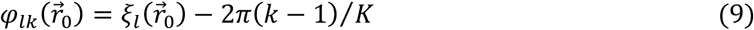

with the global phase 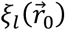 of the molecule with respect to the phase offset of the illumination patterns in direction 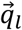 defined by:

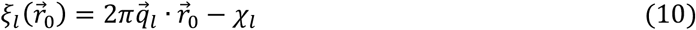

Note that the use of equidistant phases over the full 2*π* phase range ensures that the normalization condition Equation 1 is automatically satisfied. In the following, we will use this model to derive the Fisher-matrix and Cramér-Rao Lower Bound (CRLB) for the estimation of the position of the molecule.

The image formation model must be amended in case a non-zero background and/or a non-zero pixel size is taken into account. The expected photon count on pixel *j* then becomes:

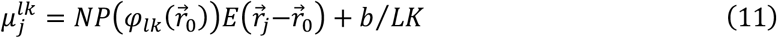

with *b* the cumulative background over *L* × *K* frames. The background is assumed to be uniform over the Region Of Interest (ROI), and constant from frame-to-frame. The integration of the PSF over the pixel area gives the factor:

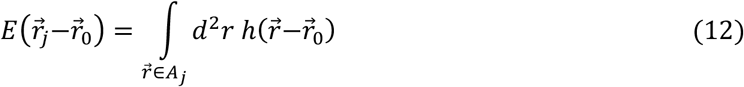

with *A_j_* the *a* × *a* sized area of pixel *j*. For the Gaussian PSF this results in:

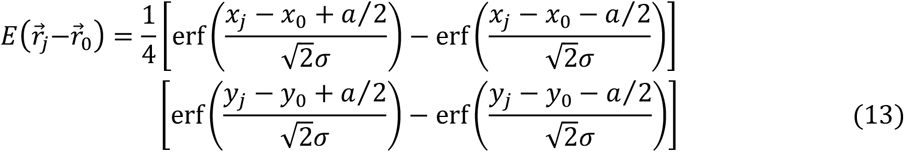

Simple analytical results for the CRLB cannot be obtained in this more general case, and we must resort to fully numerical simulations.

In the numerical analysis of our experimental results we also need to take into account that the modulation depth and the overall intensity of the illumination patterns can vary with the direction of the illumination pattern. In that case the illumination pattern for orientation *l* changes to:

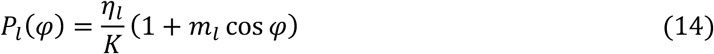

where *η_l_* is the relative intensity factor normalized as 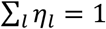, nominally *η_l_* = 1/*L*. The overall normalization condition changes to:

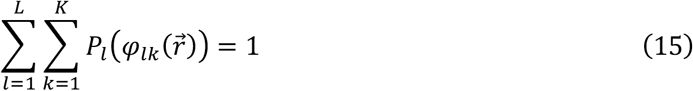

and the expected photon count on pixel *j* changes to:

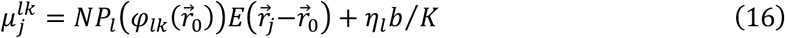

where the relative intensity also affects the background (that should scale with illumination intensity too).

### 2. Log-likelihood and derivatives

The mixed shot-noise and readout noise log-likelihood^1^ is:

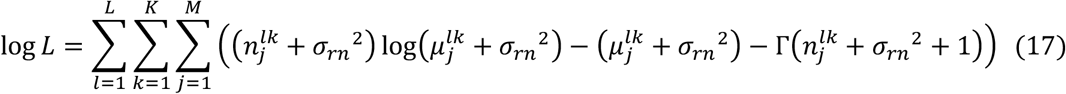

with 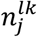 the actually detected photon count on pixel *j* for image *lk*, where *σ_rn_* is the root mean square (rms) readout noise, where 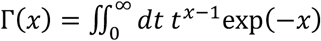 is the Gamma-function, and where the sum is over the *M* pixels of the ROI. In the numerical implementation of the MLE problem the parameters *θ* = [*x*_0_, *V*_0_, *N, b*] are estimated. This is done using the Levenberg-Marquardt routine based on the derivatives of the log-likelihood:

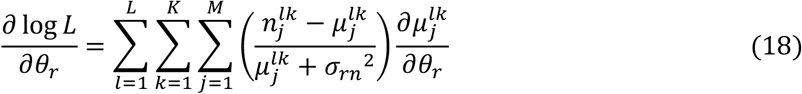

The relevant derivatives of the expected photon count on pixel *j* for image *lk* are:

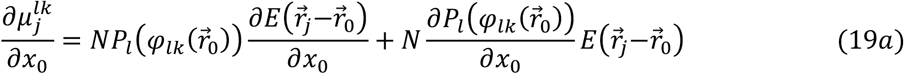

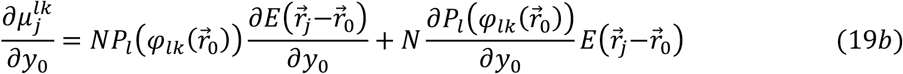

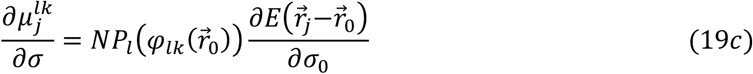

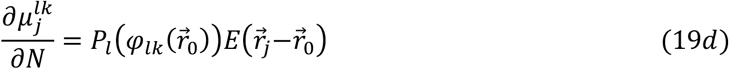

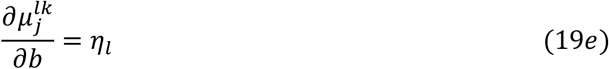

The derivatives of the Gaussian PSF term 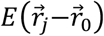 are as in Smith et al.^2^. The derivatives of the illumination pattern factor 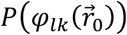 are:

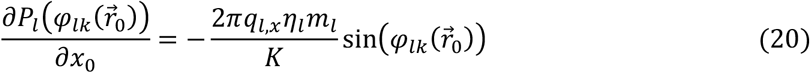

and similarly for the derivative with respect to *y*_0_. The Fisher-matrix can be computed according to:

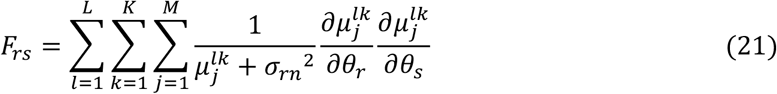

The CRLB follows from the diagonal of the inverse of the Fisher-matrix. The impact of the very small readout noise of sCMOS cameras (typically *σ_rn_* ≈ 1e) is neglected in the further analysis.

### 3. Analytical approximation Fisher-matrix and CRLB

In order to make a theoretical assessment of the expected gain in localization precision we will develop an analytical approximation to the Fisher-matrix and CRLB. This can be done in the for the case of zero background, if we ignore the non-zero pixels size, and if we neglect the finite support of the Region Of Interest (ROI). The relevant Fisher matrix elements can be written as:

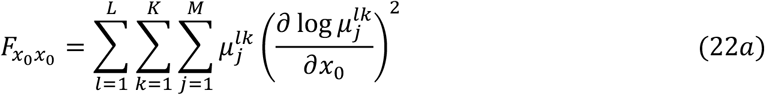

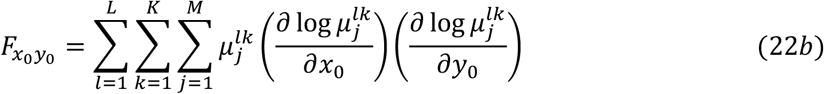

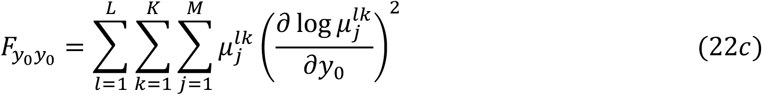

For the case with zero background and ignoring the non-zero pixel size and the direction dependence of the illumination intensity and modulation we have:

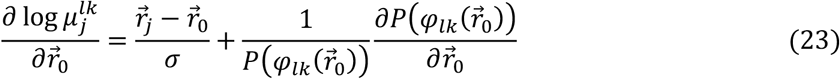

Approximating the summation over the pixels of the ROI with an integration over the entire 2D image plane results in:

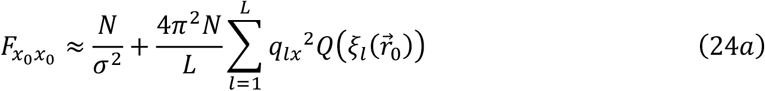

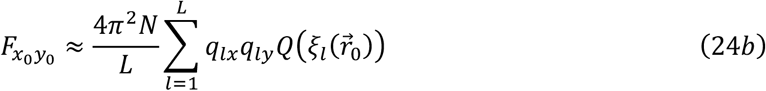

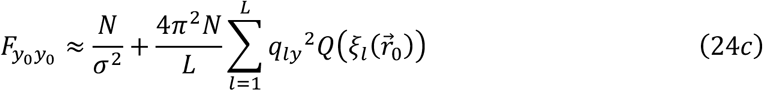

Here the function 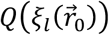 is defined by:

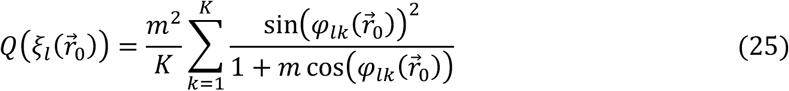

### 4. Localization precision with perfect modulation

In case of perfect modulation *m* = 1, the function 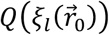 simplifies to:

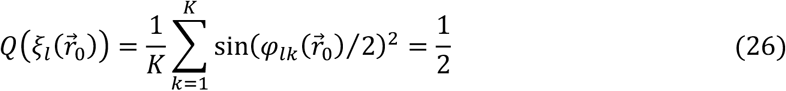

and is thus independent of position 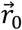 and orientation *l*. For *L* ≥ 2 (at least 2 orientations) we find:

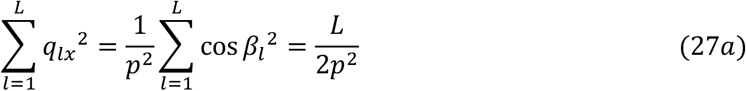

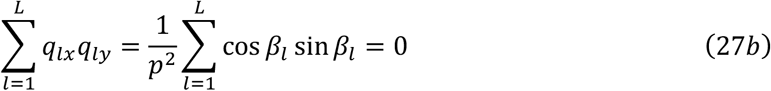

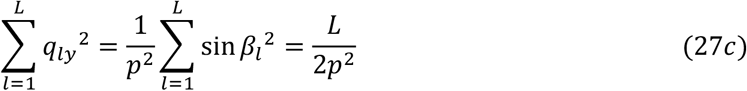

independent of the overall angle offset *β*_0_. Substituting Equations 26 and 27 in Equations 24 results in the two diagonal Fisher-matrix elements being non-zero and equal to:

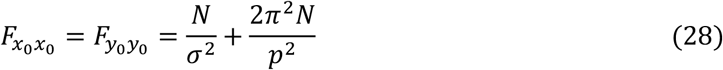

This gives an isotropic localization uncertainty:

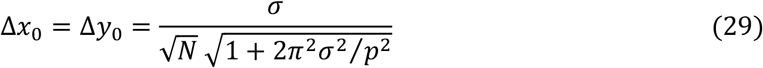

which improves over SMLM with a factor of around 2 depending on the pattern pitch *p* in relation to the spot width *σ*.

### 5. Localization precision with imperfect modulation

In case *m* < 1 the precision will become worse. Moreover, there will be a (slight) dependence on the global phase 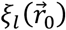 defined in Equation 10. This will make the localization precision non-uniform and anisotropic to some degree.

In order to analyse these results we will assume, for the sake of simplicity, that the illumination patterns are oriented along the the *x*-axis and *y*-axis. Following the same steps as above we then find that:

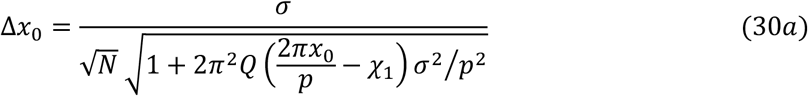

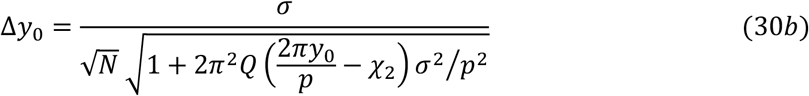

The average behaviour can be deduced by replacing the summation over the *K* phase steps in Equation 25 by an integral over all phases:

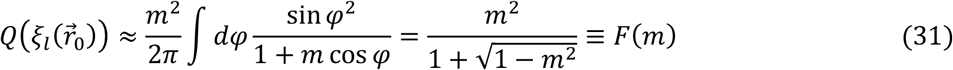

Supplementary Figure 7 shows the effect of the dependence of the localization precision on the global phase. The worst case happens when one of the phases 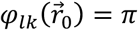 (emitter at minimum of illumination pattern for one of the frames). In case of an ideal modulation contrast *m* = 1 this is a perfectly dark fringe and this helps significantly to decrease the CRLB. In case of a non-ideal modulation contrast *m* < 1 there is no added value. For example, the improvement in CRLB for *m* = 0.95 and *K* = 3 phase steps can vary between 1.6 and 2.3, with an average of 2.1 (taking a pitch to spot width ratio *p/σ* = 2). These variations decrease when the number of phase steps is increased, making the method more robust. For example, for *m* = 0.95 and *K* =4, the improvement factor varies between 1.8 and 2.2, for *m* = 0.95 and *K* = 5, the improvement factor merely varies between 1.9 and 2.2. In practice, these variations are further mitigated by the non-zero background.

### 6. Localization precision with reduced scan range

The current results can be generalized by changing the translation range of pattern shifting from the pattern pitch *p* to a smaller range *R*, similar as in MINFLUX. So, we take pattern phases:

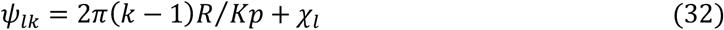

In order to enforce the normalisation condition Equation 1 (keeping the parameter *N* the number of detected photons) we must normalize the patterns with a factor:

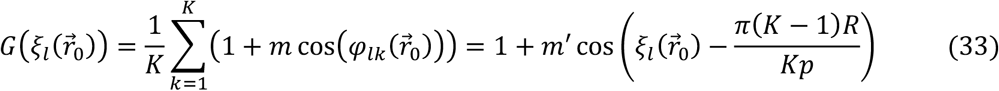

with

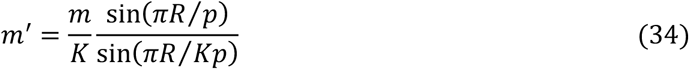

This normalization factor reaches a minimum when:

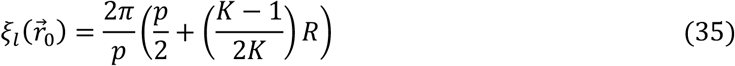

which implies that half way the scan the illumination pattern intensity minimum coincides with the molecule. The expected photon count on pixel *j* is:

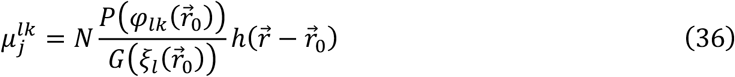

Taking the same steps as in the previous derivation results in a function 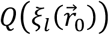 that is rather involved:

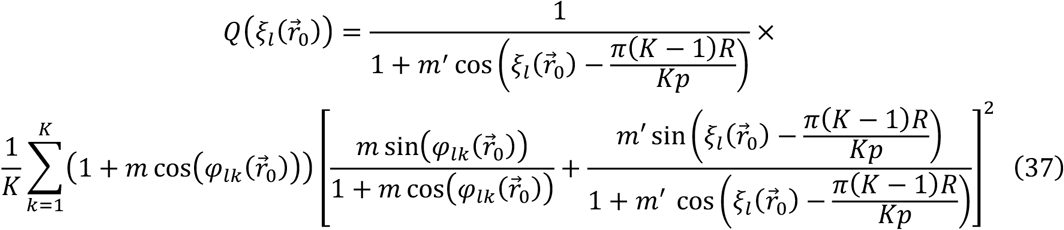

and a localization precision still given by Equations 23. For the limiting case *R* ≪ *p* and perfect modulation *m* = 1 the function 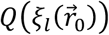 is sharply peaked around the value 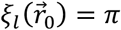, indicating that a small scan range, centred around the intensity minimum, results in a small localization precision, as in MINFLUX. At points in the FOV close to the crossing points between the intensity minimum lines of the patterns oriented along the *x*-axis and *y*-axis there will be a large improvement in precision, given that a constant photon count *N* per localization event can be achieved. Supplementary Figure 13 shows that for a perfect modulation (and zero background) an in principle unlimited improvement over SMLM can be achieved by reducing *R*, in agreement with Balzarotti et al.^3^. For an imperfect modulation, however, this is not the case. The improvement factor can reach values up to about 10, and reaches an optimum not for a global phase corresponding to the molecule being at the centre of the range of positions of the illumination pattern minimum, but rather near the edge of this range.

It is mentioned that this type of precision improvement (for a reduced phase scan range) can only be achieved for STORM type of photo-switching, where the typical number of photons per on-event is molecule specific and intensity independent, as the lower intensity near the illumination pattern minimum will reduce the number of photons per unit time, but at the same time make the molecular on-time longer. The major drawback in this case is that the longer on-time makes the on-off ratio unfavourable, more molecules per unit area will be in the on-state at any given moment in time. For PAINT type of photo-switching the on-off transition is diffusion driven, implying that the gain in localization precision per detected photon is cancelled by a reduction in the number of detected photons per on-event.

